# Theta-band phase locking during encoding leads to coordinated entorhinal-hippocampal replay

**DOI:** 10.1101/2021.10.18.464813

**Authors:** Diogo Santos-Pata, Caswell Barry, H. Freyja Ólafsdóttir

## Abstract

Precisely timed interactions between hippocampal and cortical neurons during replay epochs are thought to support memory consolidation. Indeed, research has shown replay is associated with heightened hippocampal-cortical synchrony. Yet, many caveats remain in our understanding. Namely, it remains unclear *how* this offline synchrony comes about, whether it is specific to particular behavioural states and how - if at all - it relates to learning. In this study, we sought to address these questions by analysing coordination between CA1 cells and neurons of the deep layers of the medial entorhinal cortex (dMEC) while rats learned a novel spatial task. During movement, we found a subset of dMEC cell which were particularly locked to hippocampal LFP theta-band oscillations and which were preferentially coordinated with hippocampal replay during offline periods. Further, dMEC synchrony with CA1 replay peaked ∼10ms after replay initiation in CA1, suggesting the distributed replay reflects extra-hippocampal information propagation, and was specific to ‘offline’ periods.

Finally, theta-modulated dMEC cells only became coordinated with replay after an animal’s first encounter with a novel spatial environment and then showed a striking experience-dependent increase in synchronisation with hippocampal replay trajectories, mirroring the animals’ acquisition of the novel task and coupling to the hippocampal local field. Together, these findings provide strong support for the hypothesis that synergistic hippocampal-cortical replay supports the consolidation of new memories and highlights phase locking to hippocampal theta oscillations as a potential mechanism by which such cross-structural synchrony comes about. Importantly, as CA1 phase-locking to theta is implicated in the generation of theta sequences, thought to be required for replay expression, we speculate the dMEC theta phase-locking reflects the emergence of distributed hippocampal-dMEC theta sequences and that these may support the commission of memories to long-term cortical storage.

## Introduction

The establishment of long-term episodic and spatial memories (‘memory consolidation’) is thought to be underpinned by precisely timed interactions between hippocampal and cortical circuits during periods when hippocampal cell sequences, reflecting wakeful experiences, are reactivated (‘replayed’) (Buzsaki, 1989; Marr, 1971; Olafsdottir et al., 2018; Wilson & McNaughton, 1994). Specifically, coordinated hippocampal-cortical replay is proposed to gradually establish cortical memory traces and thereby the creation of robust, persistent memories. Indeed, replay periods are associated with heightened hippocampal-cortical communication(Battaglia et al., 2004; Chrobak & Buzsaki, 1994; Maingret et al., 2016; Olafsdottir et al., 2016) and cortical cells have been found to replay synergistically with hippocampal cells(Bendor & Wilson, 2012; Ji & Wilson, 2007; Olafsdottir et al., 2016).

In previous work we showed that during hippocampal replay recorded during rest, CA1 place cells and grid cells of the deep layers of the medial entorhinal cortex (dMEC) - the principal cortical output region of the hippocampus – are functionally coordinated, depicting similar spatial positions(Olafsdottir et al., 2016). Yet, the processes during encoding that lead to this coordinated hippocampal-dMEC replay – possibly paving the way for future memory consolidation – remain largely unknown. Further, whether this cross-structural synchrony reflects learning or simply is the result of pre-existing biases in hippocampal-cortical circuits has hitherto not be investigated. Finally, recent years have seen a proliferation of studies that indicate replay may serve numerous functions in cognition including – consolidation(Ego-Stengel & Wilson, 2010; Girardeau et al., 2009; Maingret et al., 2016), planning(Jadhav et al., 2012; Pfeiffer & Foster, 2013), and memory retrieval(Carr et al., 2011; Wu et al., 2017). Importantly, the function replay serves is potentially segregated by behavioural state, with replay occurring during rest/sleep (‘offline’ periods) thought to support primarily memory consolidation(Jadhav et al., 2012; Olafsdottir et al., 2018; Olafsdottir et al., 2017). Yet, whether the dMEC participates and synchronises with all types of replay, or whether this coordination is specific to offline periods, is unknown. In this study we sought to address these questions.

To this end, we first assessed if hippocampal-dMEC replay coordination may be mediated by locking to hippocampal theta (5-12Hz) oscillations during encoding periods. Theta-band oscillations have long been implicated in hippocampal-cortical communication (Jadhav et al., 2016; Jones & Wilson, 2005; Mizuseki et al., 2009; O’Neill et al., 2013) as well as sequence-based plasticity(Drieu et al., 2018; Johnson & Redish, 2007; Muessig et al., 2019). Specifically, phase-locking of CA1 cells to the hippocampal theta-band is thought to underlie the expression of so-called ‘theta-sequences’(Guardamagna et al., 2022) which have been shown to be required for the emergence of hippocampal replay(Drieu et al., 2018). Thus, perhaps the emergence of synergistic hippocampal-dMEC replay is supported by dMEC cells locking similarly to the theta rhythm during encoding? To this end, we analysed CA1 place- and excitatory dMEC cell activity recorded in parallel while animals learned a novel spatial task and during a post-task rest period (see Olafsdottir et al. (2016)). We found a significant proportion of dMEC cells showed activity coupling that was modulated in the theta-band and whose oscillatory coupling appeared to be coordinated with the hippocampal local field potential (LFP). Moreover, we found theta-modulated dMEC cells showed strong activity and functional coordination with hippocampal replay whereas non-theta-modulated cells displayed no more coordination than expected by chance. This exclusive replay coordination of theta-modulated dMEC cells peaked 10ms after hippocampal replay, consistent with the hypothesis that the coordination reflects the projection of memories to the cortex. Importantly, these results could not be explained by differential activity rates, overlap in spatial firing fields with co-recorded CA1 cells, nor the degree or type of spatial information carried by the two dMEC groups.

Further, we found that the preferential replay coordination of theta modulated dMEC cells was modulated by behavioural state, showing significantly stronger coordination during rest periods compared to immobile periods on the track - suggesting dMEC cells synchronise selectively with replay purported to support memory consolidation. Finally, dMEC-hippocampal replay coordination showed a robust learning-related effect, emerging only after an animal’s first exposure to a novel spatial task, and then strengthening in tandem with the animal’s capability to carry out the task fluently. Importantly, this experience-dependent increase in dMEC-hippocampal synchrony was mirrored by a similar experience-dependent increase in dMEC phase-locking to the hippocampal theta rhythm.

Taken together, these findings suggest oscillatory coupling, particularly in the theta-band, between hippocampal and entorhinal cells during encoding periods may be responsible for their offline synchronisation. Further, the time lag of the dMEC-hippocampal replay synchrony, the accentuated coordination observed during offline periods and the relationship between dMEC-hippocampal replay synchrony and learning together provide compelling evidence for the hypothesis that this cross-structural synchronisation represents a hallmark of memory consolidation. Finally, as theta phase-locking is thought to be required for the generation of hippocampal theta sequences(Guardamagna et al., 2022), we speculate that the dMEC theta phase-locking may reflect the emergence of distributed hippocampal-cortical theta sequences and that this distributed temporal code may facilitate the formation of long-term memories.

## Results

We analysed co-recorded CA1 place cells and excitatory cells from the deep layers (V/VI) of the MEC (dMEC) across multiple days (Figure 1B, Fig. S1) while animals ran on a Z-shaped track for food reward (RUN), rested in a cylindrical-shaped environment outside the maze environment (REST), and finally explored an open-field environment (Figure 1A, as described previously Olafsdottir et al.(Olafsdottir et al., 2016)). Recordings took place over 2-6 days. As the identification of replay trajectories depends on cells displaying spatially confined activity, we limited our analyses to dMEC cells which fell into one of the following functional classes: grid cells(Hafting et al., 2005), head direction cells(Taube et al., 1990), conjunctive grid and head direction cells(Sargolini et al., 2006), border cells(Lever et al., 2009; Stensola et al., 2012) and other spatial cells (Fig.1C, see Materials and Methods for cell classification).

**Figure 1.**
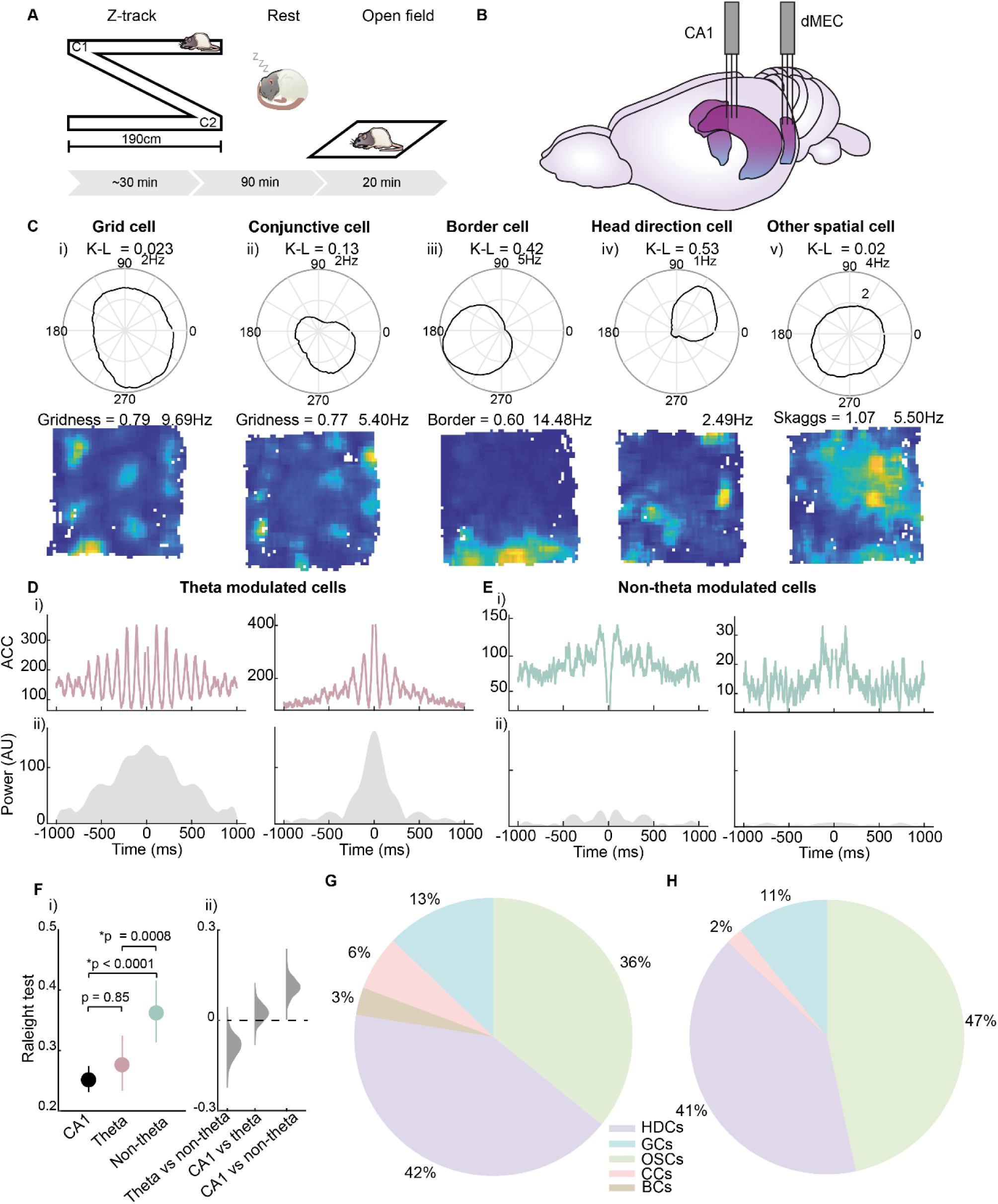
Spatial and oscillatory characterisation of dMEC cells. **A)** Schematic overview of the study. Rats shuttled back and forth between the ends of a Z-shaped track (left), and then rested for 90min in a separate environment (middle) and then completed a 20min foraging session in the open field. **B**) Two tetrode arrays were implanted in the CA1 and dMEC. **C**) Functional cell types recorded in dMEC. Top: polar plot showing directional tuning (title = direction tuning based on K-L divergence). Bottom: ratemaps showing spatial modulation, title on the left shows grid score(20) (far left, middle left), border score (21) (middle) or Skaggs information (22) (bits/spike, far right) and the peak rate (Hz), on the right. Spatial cell types from left-right: Grid cell, conjunctive grid x head direction cell, border cell, head direction cell, other spatial cell. **D**) (i) Auto-correlograms for two dMEC cells that show theta-band activity modulation. (ii) Power in the theta-band (5-12Hz) for the autocorrelogram in panel (i). **E**) Same as D but for dMEC cells that do not show significant theta-band activity modulation. **F**) (i) Mean phase locking to hippocampal theta for CA1 (black), dMEC theta-modulated (pink) and non-modulated (green) cells. Error bars show 95% CI. (ii) Kernel density function of bootstrapped difference scores. **G**) Pie chart showing contribution of different spatial cell types to the theta-modulated dMEC sub-group. **H**) Same as G) but for non-theta modulated dMEC cells. HDCs = head direction cells, GCs = grid cells, OSCs = other spatial cells, CCs = conjunctive cells, BCs = border cells.

### Theta modulation in dMEC neurons

To identify theta modulated dMEC cells we constructed an auto-correlogram for each dMEC cell spike train and assessed the power in the autocorrelogram in the theta-band (5-12Hz) relative to a broadband (20-125Hz). Cells whose theta-to-broadband ratio exceeded the 97.5^th^ percentile of its own shuffle distribution (based on random permutation of spike times) were considered theta-modulated cells (see Materials and Methods). Just over half of the dMEC cell population qualified as being theta-modulated on this measure (56.18%, 191 cells, see Figure 1D-E and Figure S2 for examples). Similarly, theta-modulated dMEC cells showed significantly stronger phase locking to hippocampal theta compared to non-theta-modulated cells (Raleigh’s test of uniformity: theta-modulated cells = 0.28 (SD = 0.31), non-theta-modulated cells = 0.36 (SD = 0.31) p = 0.008, Figure 1F). Indeed, the phase-locking to hippocampal theta exhibited by these dMEC cells was not different to that of CA1 place cells (Raleigh’s test of uniformity = 0.25 (SD = 0.30), p = 0.85, Figure 1F), suggesting that the oscillatory coupling observed in this dMEC subgroup may be synchronised with the hippocampal LFP.

Although both dMEC groups contained a mixture of different spatial cell types (see Figure 1G-H for a breakdown of the different spatial cell types for the two dMEC groups), we found some differences in the representation of individual functional cell categories between the two groups. Specifically, we found grid cells, conjunctive grid cells and border cells made up a (marginally) greater proportion of theta modulated cells compared to non-theta modulated cells (grid cells: theta-modulated = 19.25%, non-theta modulated = 12.84%, p = 0.056; conjunctive grid cells: theta-modulated = 6.42%, non-theta-modulated = 2.03%, p = 0.019; border cells: theta-modulated = 3.21%, non-theta modulated = 0%, p < 0.0001, bootstrapped proportions, Figure S3A,D-E). Conversely, other spatial cells contributed to a greater extent to the non-theta modulated subgroup (35.83% of theta-modulated and 46.62% of non-theta modulated cells, p = 0.023, Figure S3C). The proportion of head direction cells did not, however, differ between the two groups (41.71% of theta-modulated and 40.54% of non-theta modulated cells, Figure S3B). These differences are controlled for in subsequent analyses.

### Theta-modulated dMEC cells show enhanced coordinated with hippocampal replay

Next to assess if theta-band oscillatory coupling influenced the participation of dMEC cells in hippocampal replay events, we first analysed the participation of the two dMEC groups – theta-modulated and non-modulated – in candidate replay events. As this analysis does not require replay events to express coherent spatial trajectories all hippocampal MUA events submitted for replay trajectory analysis were included (‘candidate’ replay events, see Materials and Methods). Further, to control for different baseline activity levels and to assess if individual dMEC groups showed greater replay participation than expected by chance, we computed normalised replay participation scores. Namely, we permuted the timing of candidate replay events – by adding a random time shift to individual events - and determined the proportion of events each cell was active in, in these shuffled events. This process was repeated 100 times and the average shuffle score then computed. The proportion of events a cell was active in the real data was than divided by the mean of the shuffled data (a value above 1 indicates the cell is more active during events than expected by chance). We observed both dMEC subgroups were significantly more likely to participate in candidate replay events than expected by chance on this measure (theta-modulated cells = 2.10 (SD = 2.14) p < 0.0001, non-theta-modulated cells = 1.46 (SD = 1.83) p = 0.0001, based on bootstrapped participation scores, Figure 2A). However, theta-modulated cells showed significantly stronger modulation by candidate events compared to their non-theta modulated counterparts (p = 0.004, bootstrapped difference scores, Figure 2A). Further, a significantly greater proportion of theta-modulated cells showed replay participation that significantly exceeded chance levels relative to their own shuffle distribution compared to non-theta modulated cells (theta modulated cells = 46.50% (SD = 3.21), non-theta modulated cells = 30.18% (SD = 3.60), p = 0.0006, Figure 2B). Similarly, replay modulation as measured by normalised replay activity rates (i.e. average number of spikes emitted during candidate events normalised by chance) revealed the same pattern of results (mean normalised activity: theta modulated cells = 2.17 (SD = 2.44) p < 0.0001, non-theta modulated cells = 1.55 (SD = 2.07) p = 0.0002, theta vs non-theta modulated p = 0.0096; percentage of significantly modulated cells: theta modulated = 44.85% (SD = 3.18), non-theta modulated = 29.54% (SD = 3.50), theta vs non-theta modulated p = 0.001, see Figure S4A,B). Finally, a peri-stimulus-time histogram (PSTH) analysis showed that both cell groups displayed a similar temporal profile in relation to hippocampal replay events, their activity peaking ∼10ms after CA1 (see Figure 2C). Thus, both dMEC cells groups showed a lagged activity increase during hippocampal replay, but the degree of modulation was significantly stronger for the dMEC cells displaying theta-band activity rhythmicity.

**Figure 2.**
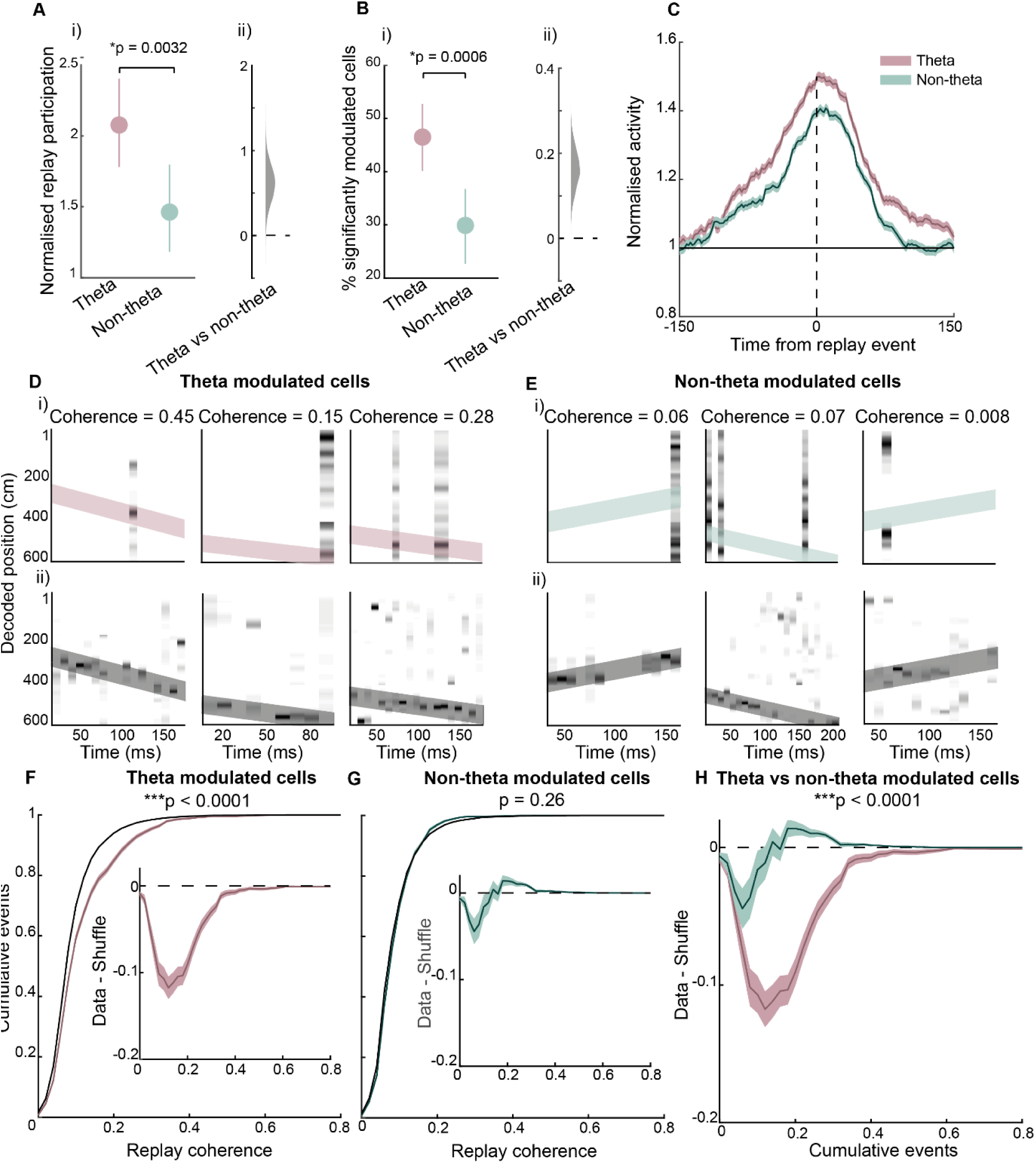
Theta-modulated dMEC cells preferentially coordinate with hippocampal replay. **A**) (i)Mean-normalised replay participation for dMEC theta-modulated and unmodulated cells. Note, a value above 1 indicate cells are more active during candidate replay than expected by chance. Error bars show 95% CIs. (ii) Kernel density of bootstrapped difference scores. **B**) (i) Mean proportion of dMEC cells that are significantly modulated by candidate replay events. Error bars show 95% CI. (ii) Kernel density for bootstrapped difference scores. **C**) Peri-stimulus-time histogram centred on the middle of candidate CA1 replay events for theta modulated and non-modulated dMEC cells. Shaded area show standard error of the mean. **D**). Representative replay trajectories with dMEC activity. (i) Bottom: position reconstruction based on CA1 spikes with best-fit line superimposed (dark grey diagonal line). Top: same as below but for dMEC theta-modulated cells, CA1 best-fit line is fitted onto dMEC decoding (pink diagonal line). Title shows dMEC-CA1 replay coherence. **E**) Same as D but for non-modulated dMEC cells. **F**). Cumulative distribution of replay coherence scores for dMEC theta-modulated cells, shaded area shows 1SD. Black line shows cumulative distribution of a cell ID shuffle. Inset: difference between the data and shuffle distribution. **G**) Same as F but for non-modulated dMEC cells. **H**) Data-shuffle distributions for the two dMEC cell types.

Next, to determine if theta-modulated dMEC cells showed stronger functional coordination with hippocampal replay, we assessed if their activity was consistent with the trajectories encoded by CA1 cells during replay trajectory events (Figure 2D-E, see Materials and Methods and Olafsdottir et al.(Olafsdottir et al., 2016)). We observed striking coordination between CA1 and dMEC theta-modulated cells, both groups expressing coherent spatial trajectories during replay events (exceeding 97.5^th^ percentile of a shuffle distribution obtained by permuting cell IDs, bootstrapped area under the curve (AUC) of data vs. shuffle: p < 0.0001, Figure 2F). Conversely, non-theta modulated dMEC cells were not coordinated with CA1 cells during replay (Figure 2G, data vs. shuffle p= 0.26) and were less coordinated than the theta-modulated cells (Figure 2H, p < 0.0001). To note, similar results were obtained when using a more stringent threshold for theta modulation (99^th^ percentile, theta-vs non-theta-modulated replay coordination: p < 0.0001, Figure S4C), a spatial firing field shuffle (theta- vs non-theta-modulated replay coordination: p < 0.0001, Figure S4D), when the number of theta and non-theta modulated cells was matched (theta-modulated vs non-modulated replay coordination after down-sampling of theta-modulated cells to match the number of non-theta-modulated cells: p < 0.0001) and when the analysis was done separately for forward and reverse replay events (forward replay: p < 0.0001, reverse replay: p < 0.0001, Figure S4E,F). Furthermore, the results were replicated for each of the three animals that individually had >10 cells in each of the two dMEC subgroups (R2335: p < 0.0001; R2336: p = 0.0008; R2337: p < 0.0001, Figure S4G-I).

### Replay coordination is not determined by activity or functional differences between theta-modulated and non-theta modulated dMEC cells

The replay coordination analysis may be sensitive to a number of functional characteristics of the dMEC cells that could differ between the theta-modulated and non-modulated subgroups. Thus, to ensure that the preferential replay coordination observed for the theta modulated sub-group could not be explained by trivial differences in activity characteristics of the two groups we carried out a number of controls. First, we assessed if the groups differed in overall activity during rest periods and the amount of spatial information carried by individual spikes (Skaggs information). The two groups displayed similar mean activity rates (theta: 0.92Hz (SD=0.90), non-theta: 0.88Hz (SD=0.88)), p = 0.36 (bootstrapped means, see Figure S5A) and spatial information (Skaggs information theta modulated cells: 1.65 (SD=0.74), non-theta modulated cells: 1.57 (SD=0.61), p = 0.85 (bootstrapped means, Figure S5C). Next, we sought to investigate if the theta modulated sub-group was more likely to have overlapping spatial firing fields with co-recorded CA1 cells than the non-theta modulated group. To this end, we computed the average spatial correlation between dMEC cells and all co-recorded CA1 cells. On average, both cell groups showed a low correlation with CA1 cells (theta modulated: r = 0.00033, non-theta modulated: r = -0.0077), which did not significantly differ from each other (p = 0.084, bootstrapped means, Figure S5B).

Further, the size of the spatial firing field of the dMEC cells could potentially influence replay coordination. Thus, we computed the average field size of the two dMEC subgroups. The spatial firing field of theta modulated cells was on average 142.17cm (SD=149.69) long whereas that of non-theta modulated cells was 171.23cm (SD=156.04), a difference we found to be statistically significant: p = 0.041. To ensure this difference could not account for their preferential coordination with CA1 replay trajectories we repeated the replay coordination analysis using a maximum field size criterion that equalised the average field size between the groups. To note, we used two different field size thresholds (150cm and 100cm), both lead to average field sizes that did not differ between the two groups (150cm control: theta modulated cells = 68.98cm (SD=31.22), non-theta modulated cells = 74.87cm (SD=31.41), p = 0.18; 100cm control: theta modulated cells = 57.64cm (SD=20.14), non-theta modulated cells = 61.28cm (SD=21.21), p = 0.26). Importantly, controlling for field size did not affect the preferential replay coordination we observed for the theta modulated sub-group (theta vs non-theta modulated coordination p < 0.0001 for both field thresholds, Figure S5D,E).

As mentioned above, some differences exist in the type of spatial cells that make up the theta modulated and non-modulated sub-groups. This could impact our results if, for example, one spatial cell category is more likely to be coordinated with hippocampal replay than another. To control for this potential confound, we carried out the replay coordination analysis separately for the different spatial cell groups. To note, as we recorded from only 15 conjunctive head direction cells (4.48%) and 6 border cells (1.79%), this analysis focused on grid (N=55, 16.42%, including conjunctive cells), head-direction (N=138, 41.19%) and other spatial cells (N=136, 40.60%). In all cases we found significantly stronger coordination with hippocampal replay for the theta-modulated sub-set of each group compared to the non-theta modulated sub-set (theta-vs non-theta-modulated replay coordination, grid cells: p < 0.0001; head direction cells: p < 0.0001; other spatial cells: p = 0.046, Figure 3A-B). Thus, the near-exclusive coordination of theta modulated dMEC cells with CA1 replay trajectories cannot be explained by some intrinsic differences between the two cell groups. Rather, we propose their replay synchronisation results from their shared oscillatory coupling with hippocampal cells.

**Figure 3.**
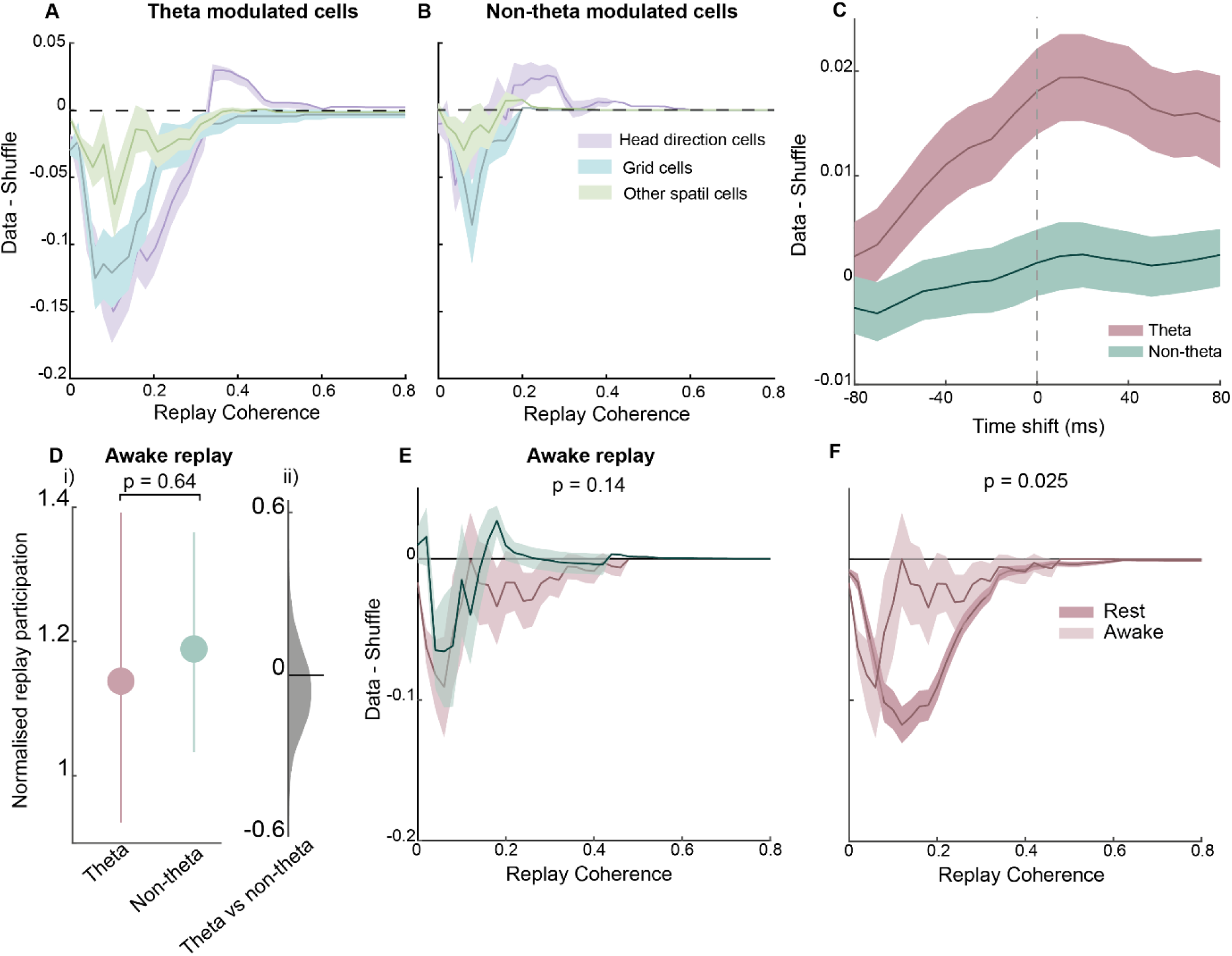
Cell-type, temporal and behavioural-state specificity of dMEC-hippocampal replay coordination. **A**) Normalised (data-shuffle) coordination between theta-modulated dMEC and hippocampal cells during replay trajectories for the different spatial cell types. **B**) Same a C) but for non-theta modulated dMEC cells. **C**) Mean dMEC-hippocampal replay coordination for different time shifts of dMEC spikes. **D**)(i) Normalised participation of theta-modulated and non-modulated dMEC cells in candidate awake hippocampal replay events. Error bars shows the 95% confidence interval. (ii) Kernel density plot of the bootstrapped difference scores. **E**) Normalised awake replay coordination for theta-modulated and non-modulated dMEC cells. Shaded area shows one standard deviation. **F**) Normalised replay coordination for theta-modulated dMEC cells during awake and rest periods

### Temporal synchronicity and behavioural state specificity of dMEC-hippocampal replay

Hippocampal-cortical coordination during replay periods is thought to support the development of complementary cortical memory traces for long-term storage(Girardeau et al., 2009; Olafsdottir et al., 2018; Wilson & McNaughton, 1994). If the dMEC-hippocampal replay coordination we observed here for theta-modulated dMEC cells indeed reflects this process, one may expect that the putative dMEC replay lags hippocampal replay - reflecting the projection of hippocampal memories to the cortex. To address this question we shifted the spike times of dMEC cells by varying amounts (+/-80ms) and computed the average coordination with hippocampal replay at each time shift. To note, to enable analysis of temporal synchrony at long time lags this analysis was limited to long replay events (>=80ms long). As expected, theta-modulated dMEC cells showed a clear peak in the coordination if we assumed the dMEC cells lagged the CA1 cells by 10ms (Figure 3C). No such peak was observed for non-theta modulated cells whose coordination with hippocampal replay remained at chance irrespective of the time shift (Figure 3C).

Thus far we have analysed hippocampal replay coordination during rest periods. Indeed, ‘offline’ replay is thought to play a privileged role in memory consolidation(Ego-Stengel & Wilson, 2010; Girardeau et al., 2009). However, replay is also known to occur during wakeful immobility periods(Diba & Buzsaki, 2007; Foster & Wilson, 2006). Importantly, this ‘online’ replay has been purported to support different functions, such as planning and decision making(Carr et al., 2011; Pfeiffer & Foster, 2013; Wu et al., 2017), although this functional distinction is still debated(Gillespie et al., 2021). Thus, we sought to investigate if dMEC cells are also coordinated with hippocampal replay that occurs while the animals are immobile on the track (<1cm/sec movement speed). In contrast to the offline analysis, we found neither dMEC cell group displayed strong recruitment to candidate replay events (normalised replay participation theta modulated = 1.14 (SD=1.37), non-theta modulated = 1.19 (SD=1.03), theta vs non-theta modulated cells: p = 0.64, Figure 3D). Indeed, only the dMEC cells that were *not* modulated by theta showed above-chance replay participation (non-theta modulated cells: p = 0.0069, theta-modulated cells: p = 0.11). With regards to coordination with hippocampal replay trajectories we also found the two cell-groups displayed similar levels of coordination (theta vs non-theta modulated cell AUC: p = 0.14, Figure 3E). Although the coordination demonstrated by the theta modulated subgroup did exceed chance levels (p = 0.017), it was significantly lower than that observed during offline replay periods (p = 0.025, Figure 3F). dMEC cells that did not exhibit theta modulated activity showed no such behavioural state-dependent change in their coordination with hippocampal replay (offline vs online replay coordination: p = 0.68, Figure S6A). To note, a similar behavioural-state dependent change in replay coordination was observed using a different movement speed threshold (3cm/sec, see Figure S6B). Thus, theta-modulated dMEC cells show accentuated coordination with hippocampal replay during offline periods - the behavioural period thought to be essential for the consolidation of new memories – strengthening the hypothesis that this coordination supports the commission of memories to long-term storage and the distinct roles of off-and on-line replay.

### dMEC-hippocampal replay synchrony requires and improves with learning

If hippocampal-dMEC replay coordination reflects the gradual commission of newly formed memories to long-term, cortical storage then one might expect the strength of hippocampal-dMEC replay synchrony to be modulated by the level of experience an animal has with a task. To address this question, we analysed separately replay coordination for three different learning periods. Namely, early (days 1-2), mid (days 3-4) and late (days 5-6). Importantly, performance on the task –measured by the number of incorrect turns at the corners of the maze - improved across these three periods (r = -0.37, p = 0.04). To note, this analysis only included the three animals that we recorded during the three learning periods so to ensure similar amount of data in each of the periods. However, the main results are replicated if all animals are included (see Figure S7A).

In terms of participation in replay, we observed theta-modulated dMEC cells were significantly more likely to participate in candidate replay events than expected by chance for all learning periods (mean replay modulation early =1.67 (SD = 2.05), p = 0.0013; mid = 2.14 (SD = 1.8); late = 2.52 (SD = 2.7), p < 0.0001, Figure 4A). Moreover, as the animal became more experienced with the task, theta-modulated dMEC cells’ participation in replay events increased, with replay participation in the late learning period being significantly higher compared to the early learning period (p = 0.028). Non-theta modulated dMEC cells, however, did not display reliable participation in replay events across the different learning periods, with only the late learning period be associated with above-chance participation (mean modulation early = 1.29 (SD = 1.85), p = 0.85; mid = 1.01 (SD = 0.82), p = 0.52; late = 2.06 (SD = 2.62) p = 0.0005, Figure 4A). In terms of coordination with replay trajectories, we found that theta-modulated dMEC cells showed marginally better coordination than expected by chance during the early learning period (p = 0.06, bootstrapped AUC), but robust coordination during the mid and late learning period (mid and late p < 0.0001, Figure 4B). Non-theta modulated dMEC cells, on the other hand, were not coordinated with replay during any learning period (early: p = 0.90, mid: p = 93, late: p = 0.11, Figure 4B). Furthermore, the theta-modulated dMEC cells showed consistently higher coordination than their non-theta modulated counterparts during all learning periods (early: p = 0.023, mid: p < 0.0001, late: p < 0.0001). However, between early and mid-learning periods we observed a significant increase in the preferential coordination of theta-modulated cells to replay compared to non-theta modulated dMEC cell (p < 0.0001, bootstrapped difference scores).

**Figure 4.**
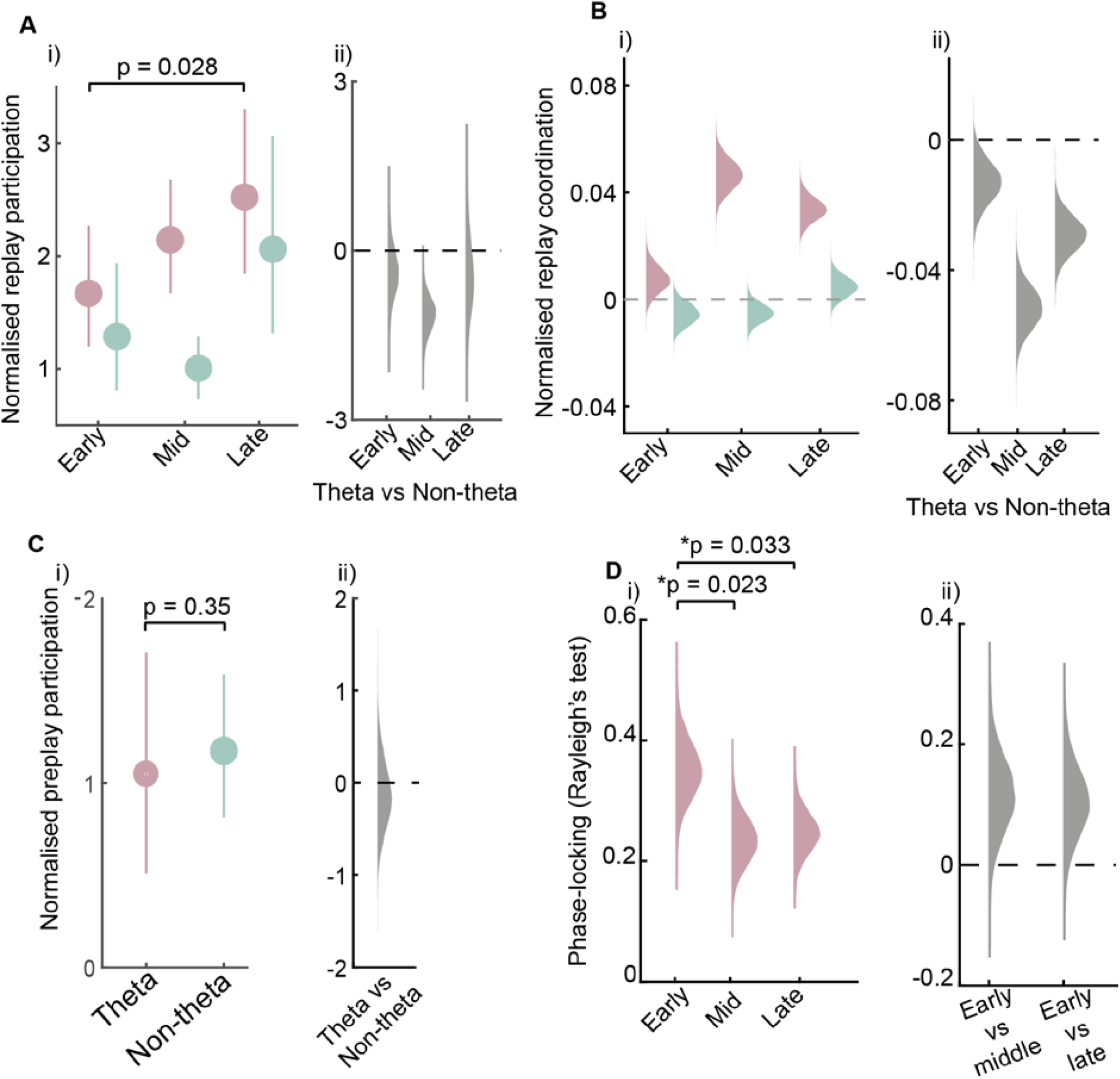
dMEC-hippocampal replay coordination of theta-modulated cells reflects learning. **A**)(i) Normalised participation of theta-modulated and non-modulated dMEC cells in candidate replay events for different learning periods. Error bars show 95% confidence interval. (ii) Kernel density function of bootstrapped difference scores (theta-modulated – non-modulated). **B**)(i) Kernel density of normalised replay coordination of theta- and non-theta modulated dMEC cells during different learning periods. (ii). Kernel density plot for bootstrapped difference scores (theta-modulated – non-modulated). **C**)(i) Normalised participation of theta- and non-theta modulated dMEC cells in candidate hippocampal preplay events. Error bars show 95% confidence interval. (ii) Kernsel density plot of difference scores. **D**)(ii) Kernsel density plot of hippocampal theta-band phase locking of theta-modulated dMEC cells during different learning periods. (ii). Kernel density plot showing difference scores for hippocampal phase locking for early vs middle and early vs late learning periods.

The foregoing analysis shows the preferential coordination of theta-modulated dMEC cells with hippocampal replay is already present in the early learning period. Thus, perhaps this preferential coordination reflects a hard-wired bias in the hippocampal-entorhinal network between CA1 and theta modulated dMEC cells (e.g. see (Dragoi & Tonegawa, 2014)). To address this question, we analysed participation in candidate (p)replay events recorded during the rest session preceding the animals’ first event encounter with the track. To note, we could not analyse coordination with replay trajectories as we did not consistently find more preplay trajectories than expected by chance during this pre sleep session. Neither theta-modulated nor non-theta modulated cells showed significant increase in their activity during the pre-sleep session (theta modulated cells = 1.05 (SD = 1.54) p = 0.53, non-theta modulated cells = 1.17, (SD = 1.05), p = 0.80, Figure 4C) and the degree of activity modulation did not differ between the two groups (p = 0.035). Thus, the preferential recruitment and synchronisation of theta-modulated dMEC cells described above emerged as an animal was exposed to and acquired a novel spatial task, and thus is likely to reflect learning-related plasticity in hippocampal-entorhinal circuits.

Finally, if the dMEC-hippocampal replay modulation for theta-modulated entorhinal cells reflects the coupling of these cells to the hippocampal theta rhythm, perhaps then the emergence of this coordination, and subsequent experience-dependent strengthening, reflects an experience-dependent increase in synchronisation of dMEC cells to the hippocampal LFP. To address this question we analysed dMEC phase-locking to the hippocampal theta rhythm during the three learning periods. Consistent with our hypothesis, we found the mean phase-locking of theta-modulated dMEC cells to increase as the animals became more experienced with the task (mean phase locking, early = 0.35 (SD = 0.36), mid = 0.24 (SD = 0.27), late = 0.25 (SD = 0.29), early vs mid: p = 0.023; early vs late: p = 0.033, Figure 4D), an effect not observed for non-theta modulated dMEC cells (early = 0.37 (SD=0.31), mid = 0.44 (SD=0.31), late = 0.33 (SD = 0.31), early vs mid: p = 0.87, early vs late: p = 0.31, Figure S7C). Indeed, phase locking to hippocampal theta by theta modulated dMEC cells became as strong as the phase locking displayed by CA1 cells during the mid and late learning period (theta modulated dMEC vs CA1 phase locking early: p = 0.013, mid: p = 0.65, late: p = 0.24). Importantly, this effect could not be explained by an increase in the number of theta-modulated dMEC cells, as the proportion of dMEC cells showing significant theta rhythmicity in their spike trains remained the same during the different learning periods (Figure S7B).

## Discussion

Influential theories posit that the formation of long-term episodic and spatial memories relies on sub-second activity synchronisation between hippocampal and cortical units during offline periods(Buzsaki, 1989; Marr, 1971; Olafsdottir et al., 2018). Yet it has hitherto remained unclear how slow, behavioural-time scale activity patterns during encoding can lead to millisecond-level, cross-regional, offline synchronisation. We show this offline coordination may be mediated by dMEC cells locking to theta oscillations during encoding periods. Specifically, we found dMEC cells that showed spike-time entrainment in the theta-band, coupled to the hippocampal local field potential, were selectively recruited to hippocampal replay events. Further, our study shows for the first time that the cross-regional synchronisation is influenced by behavioural state – being accentuated during behavioural periods thought to be the most favourable for the consolidation of new learning (i.e. rest). Finally, we found the dMEC-hippocampal replay coordination to be experience dependent, only emerging after an animal had physically explored a novel environment, and then displaying a learning-dependent increase that was mirrored by dMEC cells becoming increasingly entrained to the hippocampal theta band.

To our knowledge, this study is the first to show that hippocampal-cortical replay coordination is directly related to learning, and indeed may depend on it; cementing the hypothesis that this distributed millisecond-level synchronisation represents a hallmark of learning. Furthermore, our study highlights a candidate mechanism – namely, shared oscillatory coupling during encoding – that may be responsible for this offline synchronisation and ultimately the commission of memories to long-term storage.

The hippocampal theta rhythm has long been proposed to play a central role in extra-hippocampal communication(Buzsaki, 2010; Mizuseki et al., 2009) as well as sequence-based plasticity(Buzsaki, 2010; Drieu et al., 2018). For example, phase-locking of prefrontal cortical cells to the hippocampal theta-band has been found to be associated with enhanced re-activation during awake SWRs(Jadhav et al., 2016) and theta-nested hippocampal cell sequences have been found to be required for their later faithful reactivation during post-task rest periods(Drieu et al., 2018; Muessig et al., 2019). However, this study is the first to show that theta may also play a pivotal, potentially mechanistic, role in the establishment of offline hippocampal-cortical replay synchronisation – a process thought to lie at the heart of long-term memory formation. Given CA1 theta phase locking is thought to underlie the emergence of hippocampal theta sequences(Guardamagna et al., 2022), we hypothesise the dMEC theta phase locking we observed may reflect the emergence of distributed, hippocampal-dMEC theta sequences. As such, cross-structural sequence coupling may underlie the emergence of synchronised hippocampal-dMEC replay and thereby the commission of memories to long-term storage.

In conclusion, this study highlights and extends the role of the theta rhythm in information propagation and plasticity in hippocampal-cortical circuits, particularly in the domain of sequence plasticity. Further, finding a sub-population of cells residing in the deep-layers of the MEC may preferentially participate in hippocampal replay is consistent with the anatomical position of this subregion – i.e. it is the principle output centre of the hippocampus(Chrobak & Buzsaki, 1994; Surmeli et al., 2015) - and extends emerging theories pointing to the dMEC as a critical sub-region supporting the effective transfer of memories to the cortex(Gerlei et al., 2021).

### Acknowledgements

**Acknowledgments**: We thank Daniel Bush and Federico Stella for useful feedback on the manuscript.

## Author contributions

H.F.O and C.B. conceived of the original experiment. H.F.O performed collected the original data. D.S.P. and H.F.O. conceived, designed and performed the analyses. D.S.P. H.F.O. and C.B. wrote the manuscript.

## Data and materials availability

Data analysis code (Python) used to analyses CA1-dMEC coupling is available on Gitlab(https://gitlab.com/diogo.santos.pata/theta-coordinates-hippocampal-entorhinal-ensembles-during-wake-and-rest), code for analysing replay (Matlab) can be obtained by contacting H.F.O. The raw data can be accessed here: https://doi.org/10.5281/zenodo.5566548

## Materials and Methods

### Animals and surgery

Six male Lister Hooded rats were used in this study. All procedures were approved by the UK Home Office, subject to the restrictions and provisions contained in the Animals (Scientific Procedures) Act of 1986. All rats (330-400g at implantation) received two microdrives, each carrying eight tetrodes of twisted 17µm HM-L coated platinum iridium wire (90% and 10%, respectively; California Fine Wire), targeted to the right CA1 (ML: 2.2mm, AP: 3.8mm posterior to Bregma) and left medial entorhinal cortex (MEC) (ML = 4.5mm, AP = 0.3-0.7 anterior to the transverse sinus, angled between 8-10°). Wires were platinum plated to reduce impedance to 200-300kΩ at 1 kHz. After rats had recovered from surgery they were maintained at 90% of free-feeding weight with ad libitum access to water, and were housed individually on a 12-hr light/dark cycle, with experiments conducted during the light period.

### Recording

Screening was performed post-surgically after a 1-week recovery period. An Axona recording system (Axona Ltd.) was used to acquire the single-units and positional data (for details of the recording system and basic recording protocol see Olafsdottir et al. (2016)). The position and head direction of the animals was inferred using an overhead video camera to record the location of two light-emitting diode (LED) mounted on the animals’ head-stages (50Hz). Tetrodes were gradually advanced in 62.5um steps across days until place cells (CA1) or grid cells (MEC) were found.

### Experimental apparatus and protocol

The experiment was run during the animals’ light period to encourage quiet restfulness during the rest session. Animals ran on a Z-shaped track, elevated 75cm off the ground with 10cm wide runways. The two parallel tracks of the Z (190cm each) were connected by a diagonal section (220cm). The entire track was surrounded by plain black curtains with no distal cues. During each track session, animals were required to complete laps on the elevated Z-track, traversing each of the three tracks in order before returning in the other direction. At each end and corner, animals received a sweetened rice grain. Importantly, reward was withheld if the animal made an incorrect turn at the corners. Four animals (R2142, R2192, R2198, and R2217) were trained to run on the track for 3 days before recording commenced. For the other animals (R2242, R2335, R2336, R2337), recordings were made from the first day of exposure to the Z-track task.

Following the track session, rats were placed in the rest enclosure for 90 minutes. The rest enclosure consisted of a cylindrically shaped environment (18cm diameter, 61cm high) with a towel placed at the bottom and was located outside of the curtains which surrounded the Z-track. Animals were not able to see the surrounding room while in the rest enclosure. Prior to the experiment, rats had been familiarised with the rest enclosure for at least 7 days. Animals R2242, R2335, R2336 and R2337, were also placed in the rest enclosure for 90 minutes prior to the first Z-track session on day 1 of the experiment. Recordings from this ‘pre-rest’ session were not analysed as part of this study. Following the rest session, animals completed a 20min foraging session in an open field environment. This session was included to enable functional classification of MEC cells and was not analysed in the current study.

### Data inclusion/exclusion

Sessions recorded on days1-6 were submitted for analysis. One session was excluded as result of data loss caused by the headstages becoming disconnected from the microdrives during the rest session (R2336 day4) and one due to absence of an eeg recording (R2142, day4). In total 22 sessions were submitted for further analysis.

### Data Analysis

#### Theta modulation analysis

Theta modulation was computed for each dMEC cell individually. Specifically, each cell’s spiking activity (1ms bins) density was computed, autocorrelated (−30 to 30 sec) and convolved (20 ms). Next, theta-band modulation score was calculated by comparing the strength of theta-band (5-12Hz) against the strength of broadband (20-125Hz) and then Hilbert-transformed to obtain their respective amplitude envelope. Theta- and broad-band scores were then obtained by the theta and broad-band mean amplitude ratio within the -1 to 1 second window. The ratio between amplitude in theta and broad-band was used to theta-modulated cells. To compute statistical significance we randomly permuted the spike times of individual cells within the sessions time window. dMEC cells whose theta modulation score was above or equal the 97.5th percentile of the permutation distribution were considered to be theta modulated.

To note, only cells whose mean firing rate did not exceed 7Hz were included and whose activity fell into one of the following functional cell type categories: grid cells, head direction cells, border cells, conjunctive cells, other spatial cells. The method for each classification is described below.

#### Theta phase locking

dMEC cells were scored by their locking to ongoing hippocampal theta. To do so, we first identified the electrode in the CA1 region with strongest theta (5-12Hz) to delta (2-4 Hz) ratio. We filtered the selected CA1 channel’s signal in the theta band (finite impulse response filter, ’Hamming’ window), Hilbert-transformed it and extracted its instantaneous phase, allowing to identify the theta phase of each spike.

Only spikes elicited when the animal’s running speed was above 3cm/s were included in this analysis. Theta coherence was computed via the Rayleigh test of uniformity on each cell’s spiking theta phases. Statistical significance between cell groups was computed by bootstrapping Rayleigh test scores.

#### Hippocampal replay

Ratemaps for the Z-track were generated after first excluding areas in which the animals regularly performed non-perambulatory behaviours (e.g. eating, grooming); the final 10cm at either end of the track and 5cm around each of the two corners. Similarly, periods when the animals’ running speed was <3cm/s were also excluded. To generate ratemaps, the animals’ paths were linearised, dwell time and spikes binned into 2cm bins and smoothed with a Gaussian kernel (σ = 5bins), firing rates were calculated for each bin by dividing spike number by dwell time. Separate ratemaps were generated for runs in the outbound and inbound directions. To identify place fields, spatial bins whose rate exceeded the mean firing rate of the cell on the track were only considered. Hippocampal cells were classified as place cells if they exhibited firing greater than its mean rate for 20contiguous bins and if the peak firing rate was >1hz. These cells were submitted to further analysis. Interneurons, identified by narrow waveforms and high firing rates, were excluded from all analyses Putative replay events were identified based on the activity of hippocampal cells using a similar method to Olafsdottir et al(2016). To identify replay events, multi-unit (MU) activity from CA1 cells were binned into 1ms temporal bins and smoothed with a Guassian kernel (σ = 20ms). Periods when the MU activity exceeded the mean rate by 3 standard deviations were identified as candidate replay events. The start and end points of each candidate event were determined as the time when the MU activity fell back to the mean. Events less than 40ms long and which contained less than 15% of the recorded CA1 population (or 5 cells, which ever was greater) were rejected. Further, for the awake replay analyses we excluded all events that occurring when the animals’ average movement speed exceeded 1cm/sec. We also used a 3cm/sec movement threshold to ensure the results were not sensitive to the exact value of this threshold.

For position decoding of candidate events a Bayesian framework(Olafsdottir, et al., 2016) was used to calculate the probability of the animal’s position in each spatial bin given the observed spikes; the posterior probability matrix. Note, two posterior probability matrices were generated for each event, one for inbound runs and one for outbound runs. Spike data was divided up into 10ms temporal bins, and decoding was carried out on each bin separately.

To score the extent to which putative trajectory events represented a constant speed trajectory along the linearised Z-track we applied a line-fitting algorithm(Olafsdottir, et al., 2016). Lines were defined with a gradient (V) and intercept (c), equivalent to the velocity and starting location of the trajectory. The goodness of fit of a given line was defined as the proportion of the probability distribution that lay within 30cm of it. Specifically where P is the probability matrix:

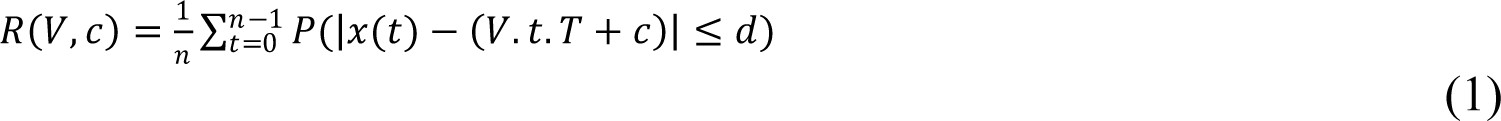

where t indexes the time bins of width and d is set to 30cm. was maximised using an exhaustive search to test all combinations of V between -50m/s and 50m/s in 0.5m/s increments (excluding slow trajectories with speeds > -2m/s and < 2m/s) and c between -15m and 21m in 0.01m increments.

To assess candidate replay events for significance we carried out a spatial field shuffle of the place cell ratemaps. Specifically, each ratemap was ‘rotated’ by shifting it relative to the track by a random number of bins drawn from a flat distribution between 1 and the length of the track minus 1 bin. The ratemap for each cell was rotated independently and in each case trailing bins were wrapped around to ensure an equal number of bins were used for each shuffle. This process was repeated 100 times for each event and for each shuffle we recalculated a goodness of fit measure (as described above). This enabled us to estimate the probability of obtaining a given event by chance. Replay trajectory events were defined as those with an individual p-value below 0.025 – to control for multiple comparisons for in- and outbound runs.

#### dMEC-hippocampal replay coordination

To analyse dMEC coordination with replay events we carried out two kinds of analyses. First, the proportion of candidate replay events (defined as those events submitted to trajectory decoding) a dMEC cell was active in was estimated for theta-modulated and non-modulated cells. To control for possible differences in baseline activity levels between the two cell types we computed the chance participation score for each individual cell by shuffling the event times of candidate events. The mean of the shuffle was then divided by the participation score in the real data. On this measure a value above 1 indicates the cell is active in more events than expected by chance. To estimate if individual cells were significantly modulated by candidate replay events the 95% confidence interval for each cell’s shuffle distribution was estimated. To assess if theta-modulated and non-modulated cell groups showed significant modulation the participation scores were bootstrapped. The difference between the two groups was then assessed for significance by computing the difference between the bootstrapped participation scores and then estimating the 95% confidence interval. To note, we also carried out an alternative activity modulation analysis where we assess if the two dMEC groups differed in how many spikes they emitted on average during candidate replay events, normalisation and statistical testing was done as before.

To investigate replay coordination between dMEC and CA1 cells we applied the same framework as we did in previous work (Olafsdottir, et al. (2016)). Namely, a bayesian decoding was done on dMEC cell spikes (to note, this analysis was done separately for theta-modulated and non-modulated dMEC cells). Hence, for each replay event we also calculated a posterior probability matrix based solely on the observed dMEC cell spikes. Rather than fitting straight-line trajectories to the dMEC cell posteriors, we compared the best-fit line from the concurrently recorded CA1 posterior. Specifically, the dMEC-CA1 cell replay coherence score was calculated using the slope and intercept parameters of the best-fit line of the accompanying CA1 event. This value we used to index replay coordination between hippocampal and dMEC cells. To estimate statistical significance of the observed coherence scores we used two different shuffling procedures. In the first instance a shuffle distribution was generated by randomly permuting the cell IDs of dMEC cells so that cells were allocated a random ratemap (from other dMEC spatial cells recorded in the session). The line fitting procedure to estimate dMEC-place cell replay coherence, described above, was re-run. To assess the statistical significance of the obtained distribution of coherence scores against the shuffle we bootstrapped the data distribution 10,000 times, computing the cumulative distribution and the corresponding area-under-the-curve (AUC, i.e. the sum of the cumulative distribution) for each bootstrap. Difference scores between each of the 10,000 AUC scores obtained from the bootstrapped data and the shuffle distribution were computed and the 95% confidence interval estimated based on these difference scores. A result was deemed statistically significant if the confidence interval did not contain 0. To compare replay coordination between theta-modulated and non-modulated dMEC cells we computed difference scores between bootstrapped AUC difference scores for the two cell types.

Second, we applied a spatial field shuffling procedure. This procedure was similar to the shuffling procedure used for hippocampal events. Specifically, each dMEC cell ratemap was shuffled by shifting it relative to the track by a random number between 10 and the length of the track minus 10 bins. The ratemap for each cell was rotated independently and trailing bins were wrapped around to ensure an equal number of bins were used for each shuffle. This process was repeated 100 times for each event. For each shuffle, the dMEC-hippocampal replay coherence score was calculated using the slope and intercept parameters of the best-fit line of the accompanying hippocampal event (unshuffled). To assess statistical significance we used an AUC test as described above.

We also repeated the analysis using a more stringent threshold for theta modulation. Namely, rather than counting dMEC cells whose theta vs broadband ratio exceeded the 97.5^th^ percentile of its own shuffle distribution as theta modulated cells we used the 99^th^ percentile instead. Similarly, we repeated the analyses after excluding dMEC cells with large spatial firing fields. For this, we used two different thresholds: 100cm and 150cm. In each case, all dMEC cells whose every spatial firing field exceeded these thresholds were excluded.

To ensure differences in replay coordination between theta-modulated and non-modulated cells could not simply be explained by differences in the number of cells belonging to each category we carried out a down-sampling analysis. Specifically, we down-sampled the theta-modulated cell population to match that of the non-modulated cell groups by removing at random cells from the theta-modulated group. For each down-sampling iteration, we obtained a bootstrapped distribution of data-shuffle AUC scores. This process we repeated a 100 times and then the average of the down-sampled bootstrapped scores was obtained. These bootstrapped difference scores were then compared to the difference scores we obtained for the non-theta modulated cell group.

To analyse replay coherence for forward and reverse replay events separately we identified replay events with a positive slope; these were classified as forward replay.

To assess the temporal synchronicity of dMEC and hippocampal replay trajectories we shifted the dMEC spike times by varying amount ranging from -80ms to +80ms. We carried out these time shifts in 10ms time bins and for each time shift we computed the average replay coordination.

#### Experience-dependent analysis

To analyse change in dMEC-CA1replay coordination as a function of experience with the task, the data was divided into three learning periods: early (days1-2), mid(days3-4) and late(days5- 6). For each learning period the mean dMEC-CA1 replay coordination or activity modulation (replay participation) was calculated and subtracted from the mean obtained from the shuffle distribution for that learning period. To estimate statistical significance the replay coherence/participation scores were bootstrapped and the mean computed for each iteration of the bootstrap. The difference between the bootstrapped data and the mean of the shuffle was then computed to assess for statistical significance. Similarly to compare changes in replay coherence between learning periods the bootstrapped difference scores, based on mean replay coherence for distinct learning periods, were computed. A similar procedure was used to assess changed in dMEC locking to hippocampal theta. Namely, the data was divided into three learning periods and the phase locking for each period bootstrapped to compute statistical significance. To compare phase locking between different learning periods, difference scores were computed for the bootstrapped data.

To note, for this analysis only data from three animals (R2335, R2336, R2337) were used as these participated in each of the learning periods. However, the main results remained the same if we included all animals.

#### Functional classification of dMEC cells

dMEC cells were classified as grid, head direction, border or other spatial cells using standard metrics, describes below. Importantly, functional classification was done based on activity recorded during open field sessions that occurred after the post-sleep session.

dMEC cells were classified as grid cells using a shuffling procedure similar to that applied elsewhere. Specifically, the hexagonal regularity of each cell was assessed using the ‘standard’ gridness measure (Hafting, 2006). The values calculated for each cell were compared with a null distribution of 100 values obtained by calculating the gridness values of data in which the cell’s spike train had been randomly permuted relative to the position of the animal by at least 30s. A cell was considered to be a grid cell and admitted to the main analysis if its standard or modified gridness value exceeded the 95th percentile of the matching null distribution.

Direction modulation was assessed by calculating the Kullback-Leibler (KL) divergence between the cell’s polar rate map and a uniform circular distribution with equal mean:

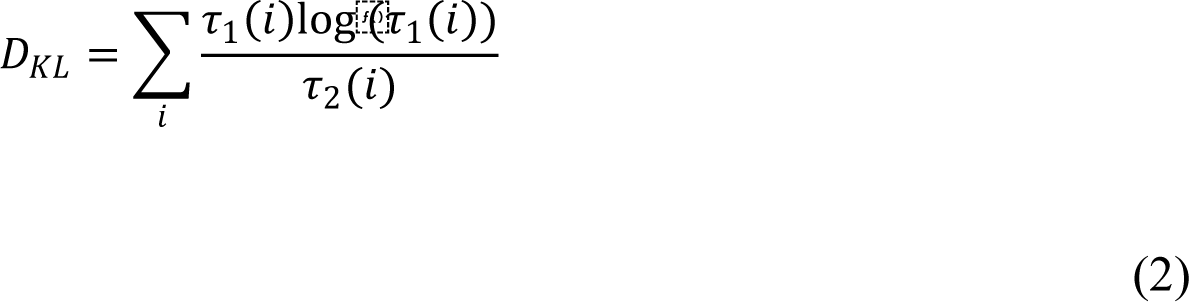

Where τ1(i) is the value in the ith bin of a polar rate map normalised to have area 1 (as a probability distribution) and τ2(i) is the ith bin of a uniform probability distribution with the same number of bins as τ1. Grid cells with KL divergence greater than 0.10 were considered to be directional.

Border score was computed as previously described (Solstad, et al., 2008). In summary, each cell’s firing fields were estimated by identifying groups of continuous spatial bins (bin size = 2cm) where the firing rate was above 30% of the cell’s peak firing rate and smaller than 70% of the arena’s area. Next, a border score (in the -1 to 1 range) was computed for each boundary individually by computing the relation between the firing field’s extent and mean distance to the wall. As in Solstad et al. (2008), cells with a border score above 0.5 were considered border cells.

Spatial modulation was assessed using Skagg’s information (Skaggs, et al. 1993). Cells whose Skaggs information (bits/spike) exceed 1 were considered as spatial cells. To note spatial cells were those cells that were not classified as any of the other spatial cell types described above.

To compare differences in representation of the distinct functional cell types for the theta-modulated and non-modulated cell groups we bootstrapped the proportions of each cell type for the two groups and computed difference scores.

#### dMEC-CA1 field overlap

In order to account for potential confounds derived from similar spatial tuning between dMEC and CA1 cells, we quantified the overlap of each cell pair rate maps during Z-track for each running direction (in- and out-bound). Only spatial bins with positive rates were included. dMEC-CA1 field overlap was scored by computing the correlation coefficient (Pearson-r test) between the two cells rate maps. To assess if theta-modulated dMEC cells displayed greater spatial firing field overlap with CA1 cells than non-theta-modulated cells we bootstrapped the ratemap correlation distributions for each group and then computed difference scores.

#### Performance analysis

To estimate performance on the Z-track, we counted the number of incorrect turns on the corners of the track and divided this by the total number of laps completed by an animal in a session. This measure gives the proportion of incorrect corner turns – a lower number indicate better performance. To assess if performance improved with experience we correlated proportion of incorrect turns with the number of days of experience using a Pearson correlation. Given we assumed an improvement with experience we report the statistics of a one-sided test.

#### Histology

Rats were anaesthetised (4% isoflurane and 4L/min O2), injected intra-peritoneal with an overdose of Euthatal (sodium pentobarbital) after which they were transcardially perfused with saline followed by a 4% paraformaldehyde solution (PFA). Brains were carefully removed and stored in PFA which was exchanged for a 4% PFA solution in PBS (phosphate buffered saline) with 20% sucrose 2-3 days prior to sectioning. Subsequently, 40-50μm frozen coronal sections were cut using a cryostat, mounted on gelatine-coated glass slides and stained with cresyl violet. Images of the sections were acquired using an Olympus microscope, Xli digital camera (XL Imaging Ltd.). Sections in which clear tracks from tetrode bundles could be seen were used to confirm CA1 recording locations.

**Figure S1.**
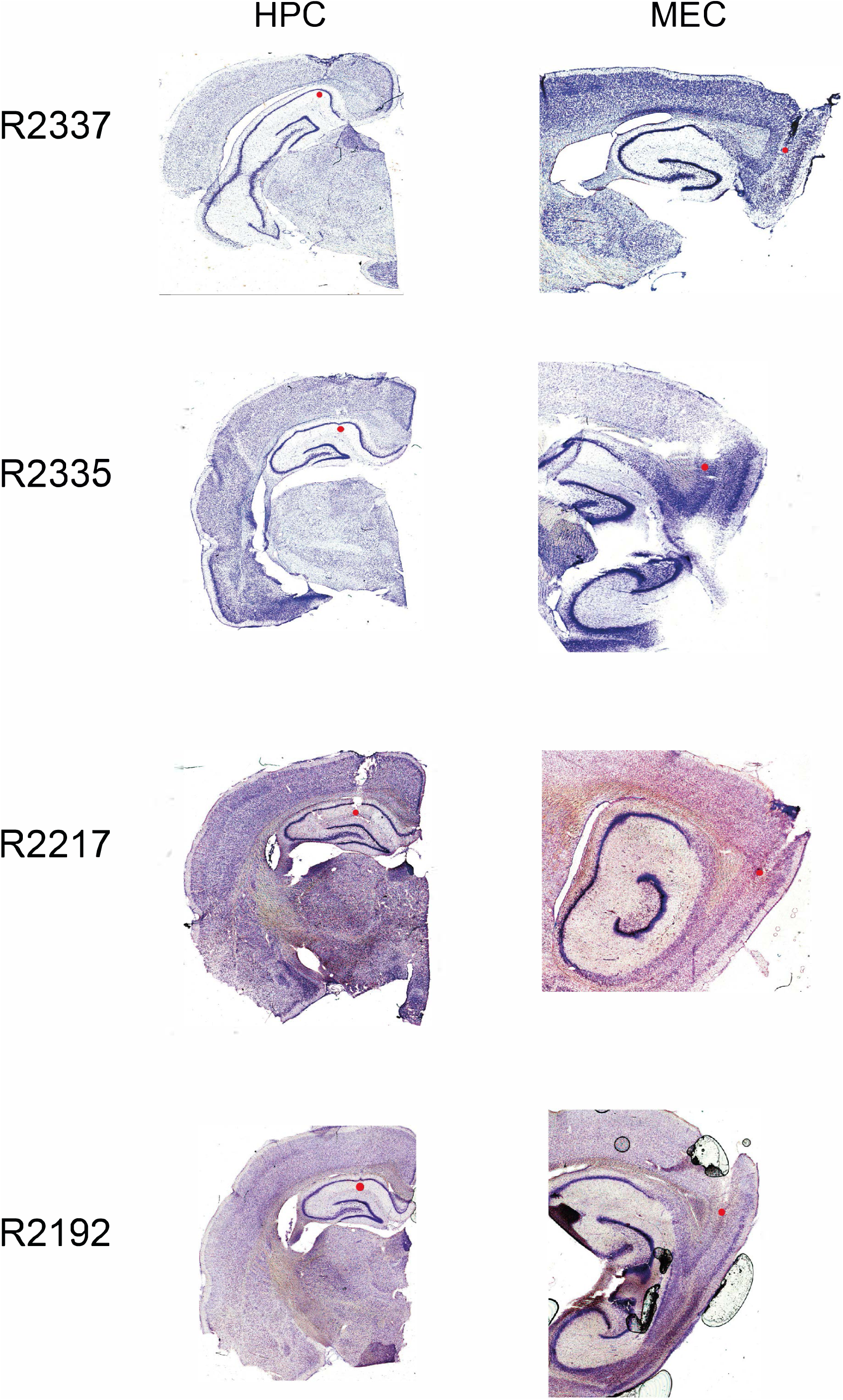
Tetrode locations. Four representative examples of Cresyl violet stained tetrode tracts from coronal (Hippocampus, left) and sagittal (MEC, right) sections. Red circle indicates the recording location for data included in this study. Left column shows rat ID.

**Figure S2.**
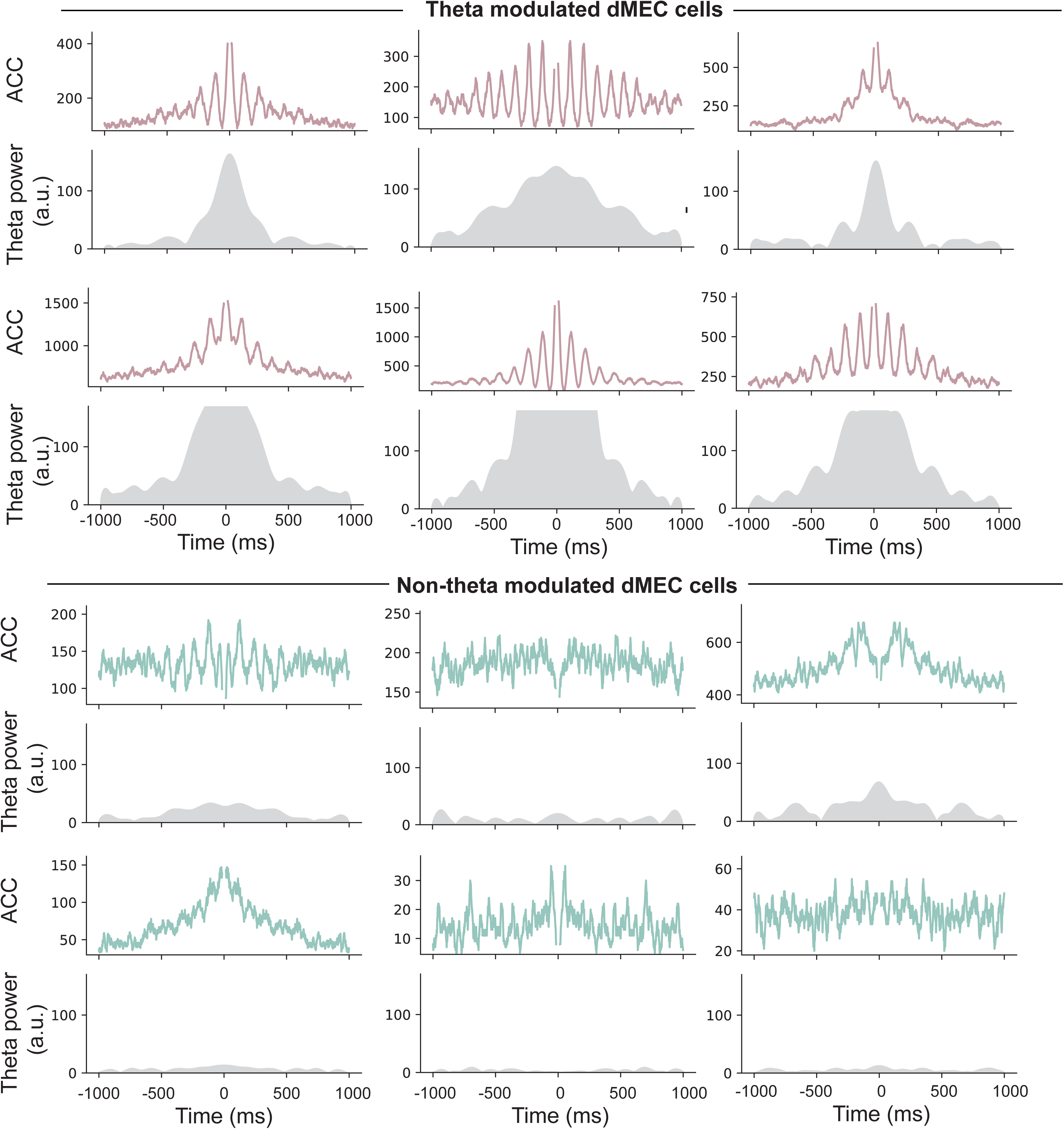
Theta-band rhythmicity in dMEC cells. Top panels: autocorrelograms for theta-modulated (pink) and non-modulated (green) dMEC cells. Bottom panel: power in the theta-band (5-12Hz) in the autocorrelogram.

**Figure S3.**
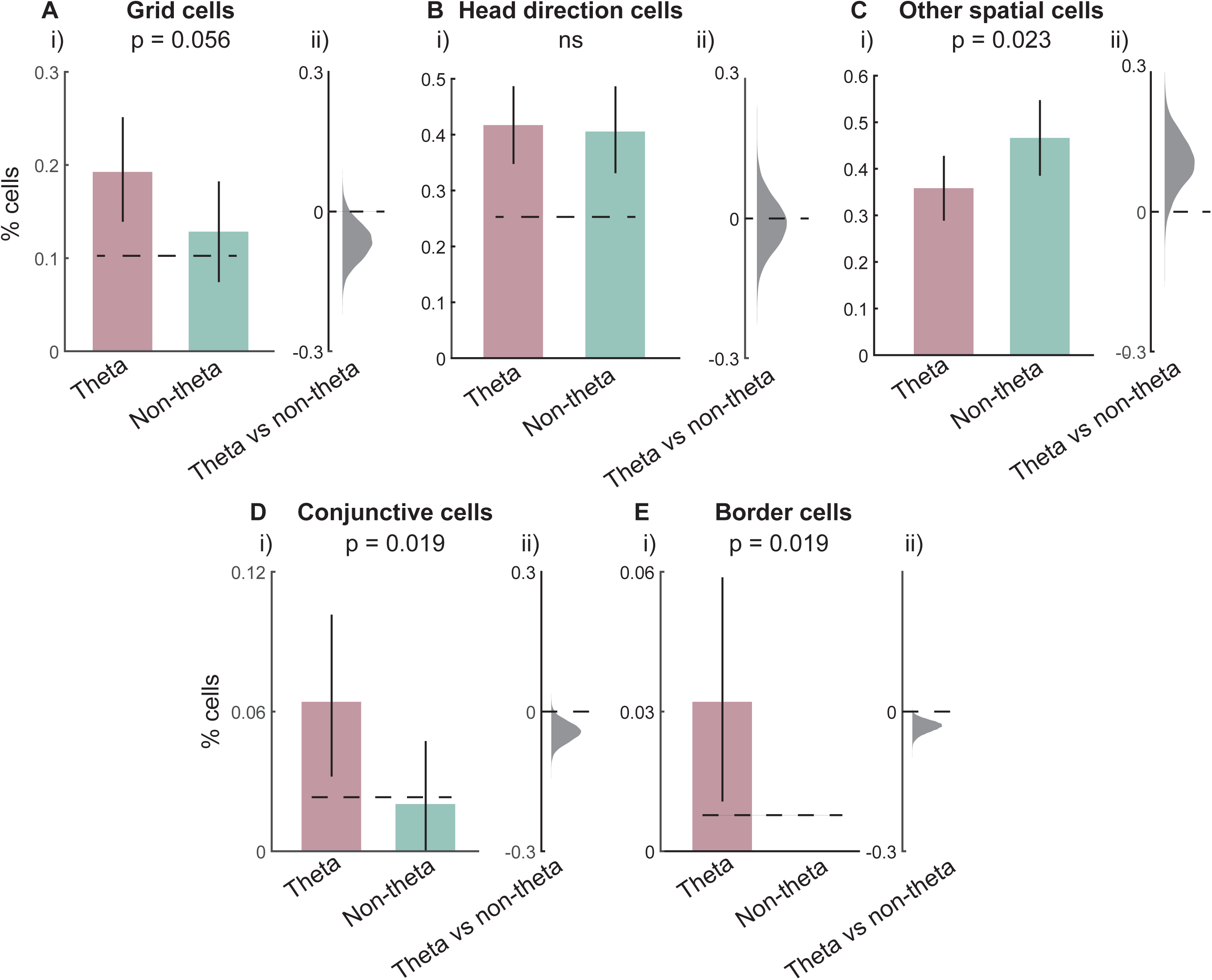
Representation of distinct functional cell types in the dMEC. **A**) i) Proportion of dMEC theta-modulated (pink) and non-modulated (green) dMEC cells that qualify as grid cells. ii) Kernel density of bootstrapped difference scores. **B-E**) Same as A but for head direction, other spatial, conjunctive and border cells. Error bars shows 95% CI.

**Figure S4.**
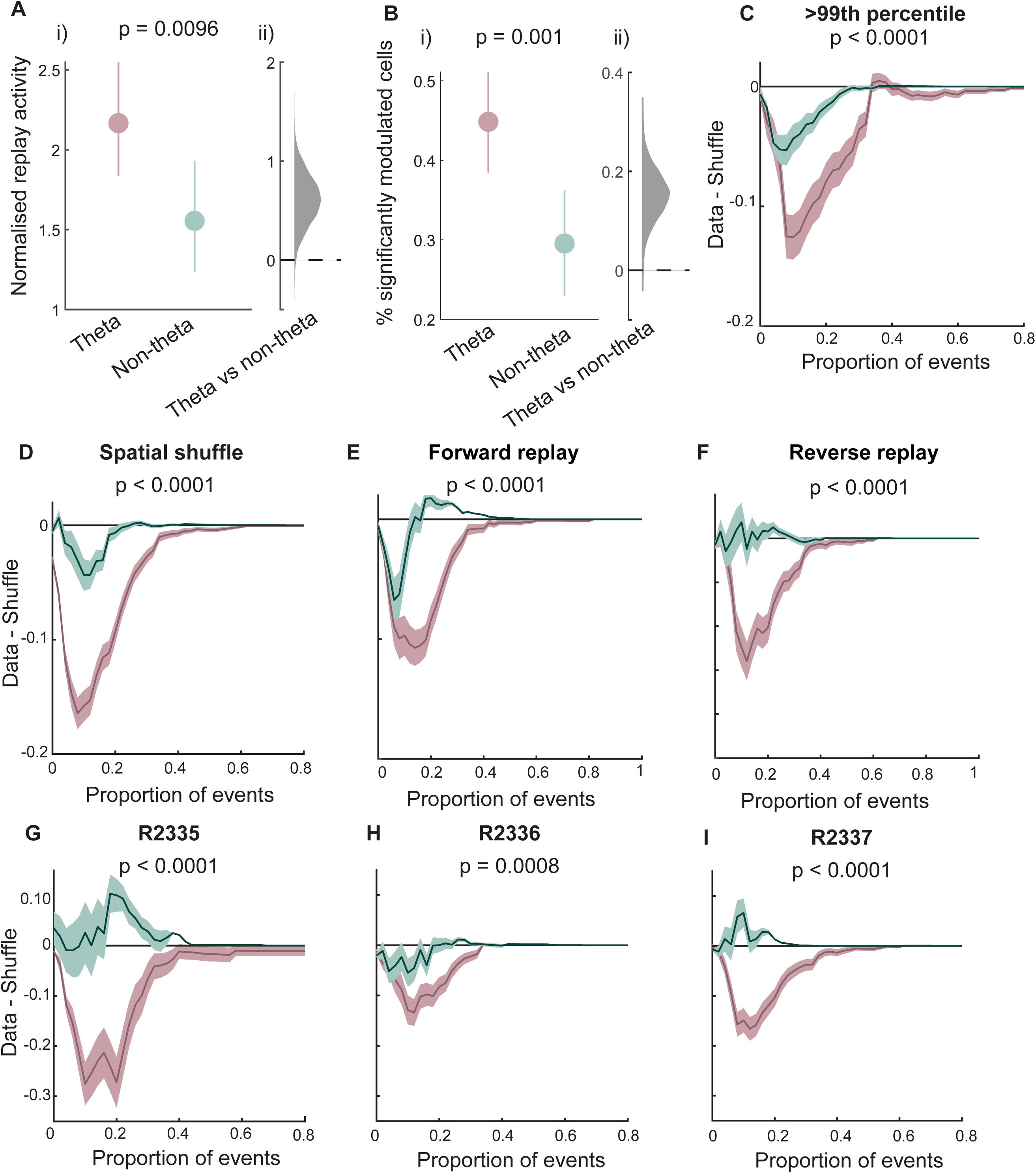
Hippocampal-dMEC replay coordination: Alternative and control analysis. **A)** i) Mean normalised activity modulation of dMEC theta modulated (pink) and non-modulated (green) cells. Error bars show 95% CI. ii) Kernel density of bootstrapped difference scores. **B**) i) Proportion of dMEC theta modulated and non-modulated cells that are significantly modulated by replay events, using meeasure from (A). Error bars show 95% CI. ii) Bootstrapped difference scores. **C**) Normalised (data-shuffle) replay coordination between hippocampal and theta and non-theta modulated dMEC cells using a more stringent threshold for theta modulation (99th percentile). Sahded area shows 1SD of bootstrapped data. **D-I)** Same as C but using an alternative spatial field shuffle (D), limiting analysis to forward (E) or reverse (F) replay events and repeating the analysis for individual animals (G-I).

**Figure S5.**
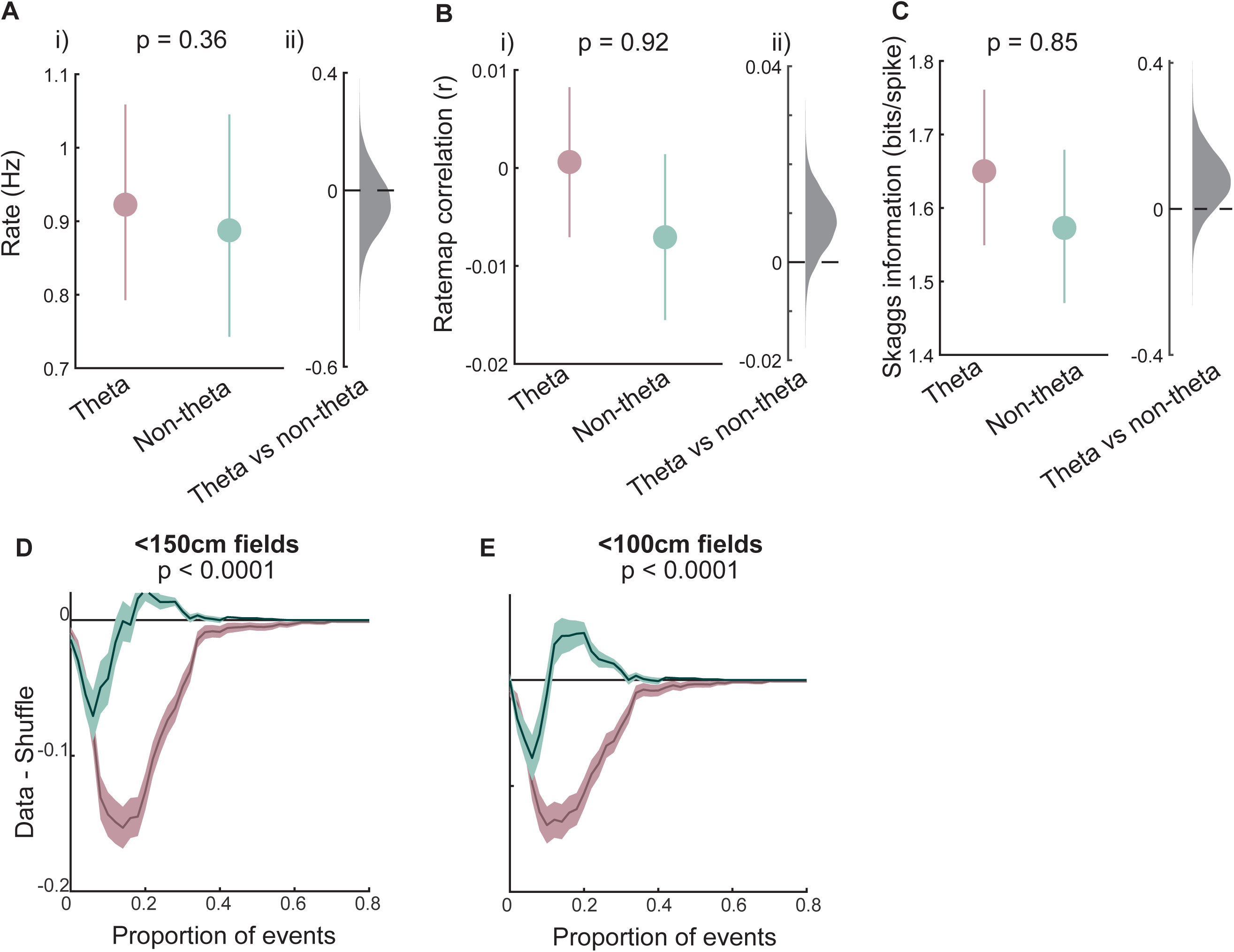
Preferential coordination of theta modulated dMEC cells with hippocampal replay trajectories is not confounded by functional and activity differences between theta-modulated and non-modulated dMEC cells. **A)** (i) Mean firing rate of theta modulated (pink) and non-theta modulated dMEC cells. Error bars show 95% CI. (ii) Kernel density of bootstrapped difference scores. **B-C)** Same as A but showing average ratemap correlations between dMEC and CA1 cells (B) and Skaggs information (C). **D)** Normalised (data-shuffle) dMEC-hippocampal replay coordination after removing dMEC cells with large (>=150cm) spatial firing fields. Shaded area shows 1SD of bootstrappe data. **E)** Same as D but using an alternative field size threshold (100cm).

**Figure S6.**
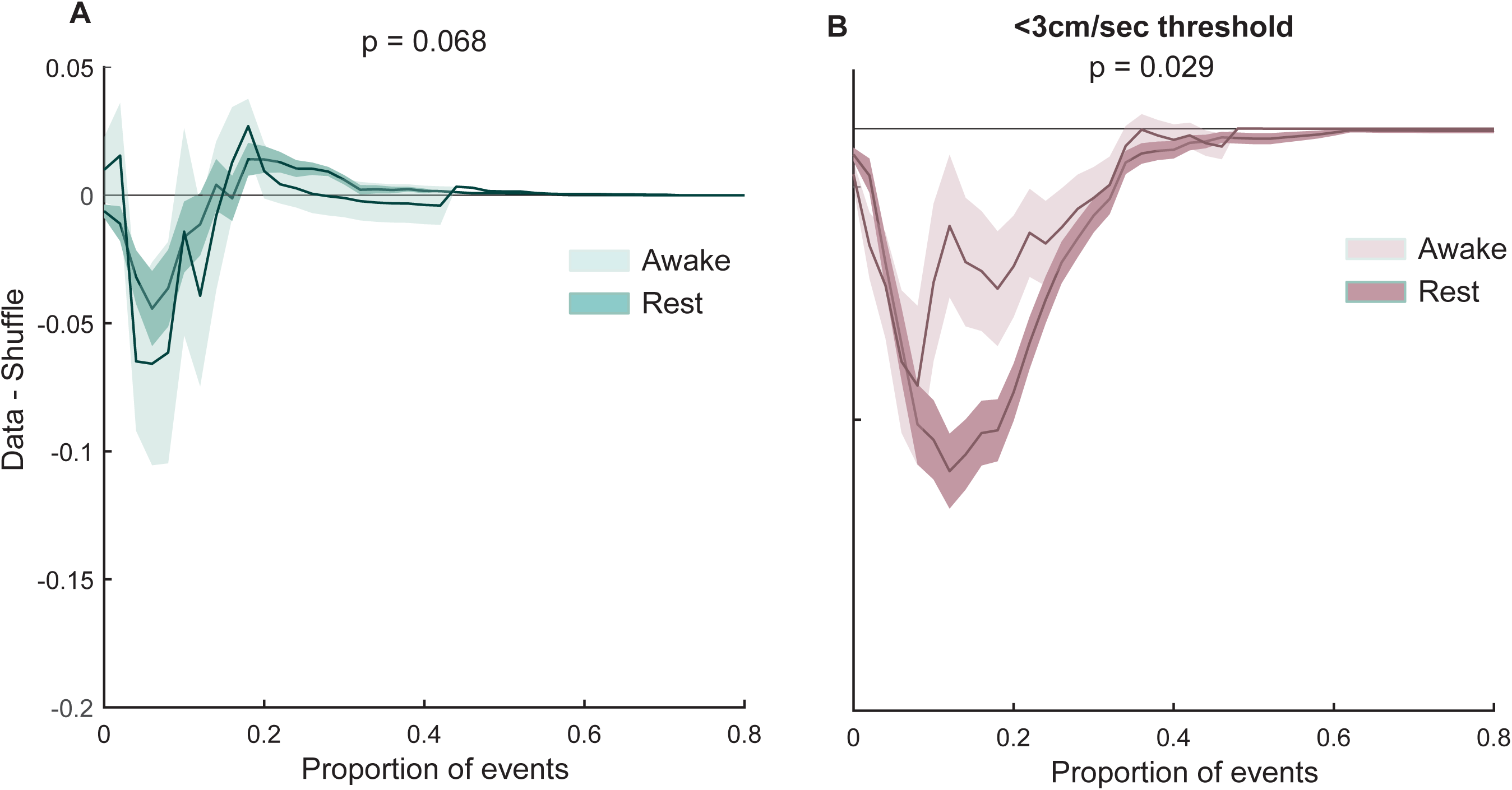
Behavioural-state dependence of dMEC-hippocampal replay coordination. **A**) Normalised (data-shuffle) replay coordination for non-theta modulated dMEC cells during awake (light green) and rest (dark green) periods. **B)** Normalised replay coordination for theta modulated dMEC cells during awake (light pink) and rest (dark pink) periods. Shaded are shows 1SD of bootstrapped data.

**Figure S7.**
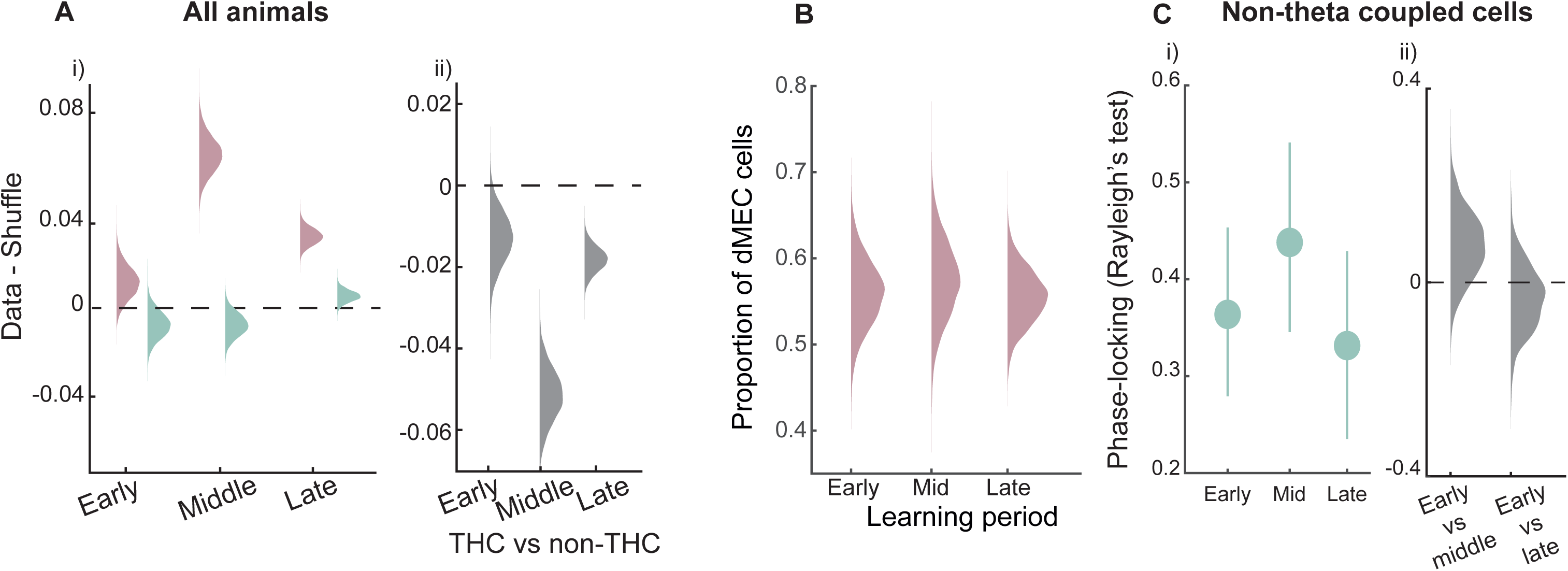
Experience dependent changes in dMEC-Hippocampal replay coordination and dMEC theta modulation. **A)** (i) Normalised (data-shuffle) replay coordination for theta-modulated (pink) and non-modulated (green) dMEC cells during distinct learning periods. Note, analysis includes all animals. (ii) Kernel density of bootstrapped difference scores between theta and non-theta modulated dMEC cells for individual learning periods. **B**) Kernel density of bootstrapped proportion of theta modulated dMEC cells during different learning periods. **C**) (i) Mean phase locking of non-theta modulated dMEC cells to hippocampal theta for different learning periods. Error bars show 95% CI. (ii) Kernel density of bootstrapped difference scores between early and mid and early and late learning periods.

## References

1. Battaglia, F. P., Sutherland, G. R., & McNaughton, B. L. (2004). Hippocampal sharp wave bursts coincide with neocortical “up-state” transitions. Learn Mem, 11(6), 697–704. https://doi.org/10.1101/lm.73504

2. Bendor, D., & Wilson, M. A. (2012). Biasing the content of hippocampal replay during sleep. Nat Neurosci, 15(10), 1439–1444. https://doi.org/10.1038/nn.3203

3. Buzsaki, G. (1989). Two-stage model of memory trace formation: a role for “noisy” brain states. Neuroscience, 31(3), 551–570. https://www.ncbi.nlm.nih.gov/pubmed2687720

4. Buzsaki, G. (2010). Neural syntax: cell assemblies, synapsembles, and readers. Neuron, 68(3), 362–385. https://doi.org/10.1016/j.neuron.2010.09.023

5. Carr, M. F., Jadhav, S. P., & Frank, L. M. (2011). Hippocampal replay in the awake state: a potential substrate for memory consolidation and retrieval. Nat Neurosci, 14(2), 147–153. https://doi.org/10.1038/nn.2732

6. Chrobak, J. J., & Buzsaki, G. (1994). Selective activation of deep layer (V-VI) retrohippocampal cortical neurons during hippocampal sharp waves in the behaving rat. J Neurosci, 14(10), 6160–6170. https://www.ncbi.nlm.nih.gov/pubmed/7931570

7. Diba, K., & Buzsaki, G. (2007). Forward and reverse hippocampal place-cell sequences during ripples. Nat Neurosci, 10(10), 1241–1242. https://doi.org/10.1038/nn1961

8. Dragoi, G., & Tonegawa, S. (2014). Selection of preconfigured cell assemblies for representation of novel spatial experiences. Philos Trans R Soc Lond B Biol Sci, 369(1635), 20120522. https://doi.org/10.1098/rstb.2012.0522

9. Drieu, C., Todorova, R., & Zugaro, M. (2018). Nested sequences of hippocampal assemblies during behavior support subsequent sleep replay. Science, 362(6415), 675–679. https://doi.org/10.1126/science.aat2952

10. Ego-Stengel, V., & Wilson, M. A. (2010). Disruption of ripple-associated hippocampal activity during rest impairs spatial learning in the rat. Hippocampus, 20(1), 1–10. https://doi.org/10.1002/hipo.20707

11. Foster, D. J., & Wilson, M. A. (2006). Reverse replay of behavioural sequences in hippocampal place cells during the awake state. Nature, 440(7084), 680–683. https://doi.org/10.1038/nature04587

12. Gerlei, K. Z., Brown, C. M., Surmeli, G., & Nolan, M. F. (2021). Deep entorhinal cortex: from circuit organization to spatial cognition and memory. Trends Neurosci. https://doi.org/10.1016/j.tins.2021.08.003

13. Gillespie, A. K., Astudillo Maya, D. A., Denovellis, E. L., Liu, D. F., Kastner, D. B., Coulter, M. E., Roumis, D. K., Eden, U. T., & Frank, L. M. (2021). Hippocampal replay reflects specific past experiences rather than a plan for subsequent choice. Neuron, 109(19), 3149–3163 e3146. https://doi.org/10.1016/j.neuron.2021.07.029

14. Girardeau, G., Benchenane, K., Wiener, S. I., Buzsaki, G., & Zugaro, M. B. (2009). Selective suppression of hippocampal ripples impairs spatial memory. Nat Neurosci, 12(10), 1222–1223. https://doi.org/10.1038/nn.2384

15. Guardamagna, M., Stella, F., & Battaglia, F. P. (2022). Heterogeneity of network and coding states in CA1. bioRxiv, 2021.2012.2022.473863. https://doi.org/10.1101/2021.12.22.473863

16. Hafting, T., Fyhn, M., Molden, S., Moser, M. B., & Moser, E. I. (2005). Microstructure of a spatial map in the entorhinal cortex. Nature, 436(7052), 801–806. https://doi.org/10.1038/nature03721

17. Jadhav, S. P., Kemere, C., German, P. W., & Frank, L. M. (2012). Awake hippocampal sharp-wave ripples support spatial memory. Science, 336(6087), 1454–1458. https://doi.org/10.1126/science.1217230

18. Jadhav, S. P., Rothschild, G., Roumis, D. K., & Frank, L. M. (2016). Coordinated Excitation and Inhibition of Prefrontal Ensembles during Awake Hippocampal Sharp-Wave Ripple Events. Neuron, 90(1), 113–127. https://doi.org/10.1016/j.neuron.2016.02.010

19. Ji, D., & Wilson, M. A. (2007). Coordinated memory replay in the visual cortex and hippocampus during sleep. Nat Neurosci, 10(1), 100–107. https://doi.org/10.1038/nn1825

20. Johnson, A., & Redish, A. D. (2007). Neural ensembles in CA3 transiently encode paths forward of the animal at a decision point. J Neurosci, 27(45), 12176–12189. https://doi.org/10.1523/JNEUROSCI.3761-07.2007

21. Jones, M. W., & Wilson, M. A. (2005). Theta rhythms coordinate hippocampal-prefrontal interactions in a spatial memory task. PLoS Biol, 3(12), e402. https://doi.org/10.1371/journal.pbio.0030402

22. Lever, C., Burton, S., Jeewajee, A., O’Keefe, J., & Burgess, N. (2009). Boundary vector cells in the subiculum of the hippocampal formation. J Neurosci, 29(31), 9771–9777. https://doi.org/10.1523/JNEUROSCI.1319-09.2009

23. Maingret, N., Girardeau, G., Todorova, R., Goutierre, M., & Zugaro, M. (2016). Hippocampo-cortical coupling mediates memory consolidation during sleep. Nat Neurosci, 19(7), 959–964. https://doi.org/10.1038/nn.4304

24. Marr, D. (1971). Simple memory: a theory for archicortex. Philos Trans R Soc Lond B Biol Sci, 262(841), 23–81. https://www.ncbi.nlm.nih.gov/pubmed/4399412

25. Mizuseki, K., Sirota, A., Pastalkova, E., & Buzsaki, G. (2009). Theta oscillations provide temporal windows for local circuit computation in the entorhinal-hippocampal loop. Neuron, 64(2), 267–280. https://doi.org/10.1016/j.neuron.2009.08.037

26. Muessig, L., Lasek, M., Varsavsky, I., Cacucci, F., & Wills, T. J. (2019). Coordinated Emergence of Hippocampal Replay and Theta Sequences during Post-natal Development. Curr Biol, 29(5), 834–840 e834. https://doi.org/10.1016/j.cub.2019.01.005

27. O’Neill, P. K., Gordon, J. A., & Sigurdsson, T. (2013). Theta oscillations in the medial prefrontal cortex are modulated by spatial working memory and synchronize with the hippocampus through its ventral subregion. J Neurosci, 33(35), 14211–14224. https://doi.org/10.1523/JNEUROSCI.2378-13.2013

28. Olafsdottir, H. F., Bush, D., & Barry, C. (2018). The Role of Hippocampal Replay in Memory and Planning. Curr Biol, 28(1), R37–R50. https://doi.org/10.1016/j.cub.2017.10.073

29. Olafsdottir, H. F., Carpenter, F., & Barry, C. (2016). Coordinated grid and place cell replay during rest. Nat Neurosci, 19(6), 792–794. https://doi.org/10.1038/nn.4291

30. Olafsdottir, H. F., Carpenter, F., & Barry, C. (2017). Task Demands Predict a Dynamic Switch in the Content of Awake Hippocampal Replay. Neuron, 96(4), 925–935 e926. https://doi.org/10.1016/j.neuron.2017.09.035

31. Pfeiffer, B. E., & Foster, D. J. (2013). Hippocampal place-cell sequences depict future paths to remembered goals. Nature, 497(7447), 74–79. https://doi.org/10.1038/nature12112

32. Sargolini, F., Fyhn, M., Hafting, T., McNaughton, B. L., Witter, M. P., Moser, M. B., & Moser, E. I. (2006). Conjunctive representation of position, direction, and velocity in entorhinal cortex. Science, 312(5774), 758–762. https://doi.org/10.1126/science.1125572

33. Stensola, H., Stensola, T., Solstad, T., Froland, K., Moser, M. B., & Moser, E. I. (2012). The entorhinal grid map is discretized. Nature, 492(7427), 72–78. https://doi.org/10.1038/nature11649

34. Surmeli, G., Marcu, D. C., McClure, C., Garden, D. L. F., Pastoll, H., & Nolan, M. F. (2015). Molecularly Defined Circuitry Reveals Input-Output Segregation in Deep Layers of the Medial Entorhinal Cortex. Neuron, 88(5), 1040–1053. https://doi.org/10.1016/j.neuron.2015.10.041

35. Taube, J. S., Muller, R. U., & Ranck, J. B., Jr. (1990). Head-direction cells recorded from the postsubiculum in freely moving rats. II. Effects of environmental manipulations. J Neurosci, 10(2), 436–447. https://www.ncbi.nlm.nih.gov/pubmed/2303852

36. Wilson, M. A., & McNaughton, B. L. (1994). Reactivation of hippocampal ensemble memories during sleep. Science, 265(5172), 676–679. https://www.ncbi.nlm.nih.gov/pubmed/8036517

37. Wu, C. T., Haggerty, D., Kemere, C., & Ji, D. (2017). Hippocampal awake replay in fear memory retrieval. Nat Neurosci, 20(4), 571–580. https://doi.org/10.1038/nn.4507

