## Supplementary material for "Theta-band phase locking during encoding leads to coordinated entorhinal-hippocampal replay": S6

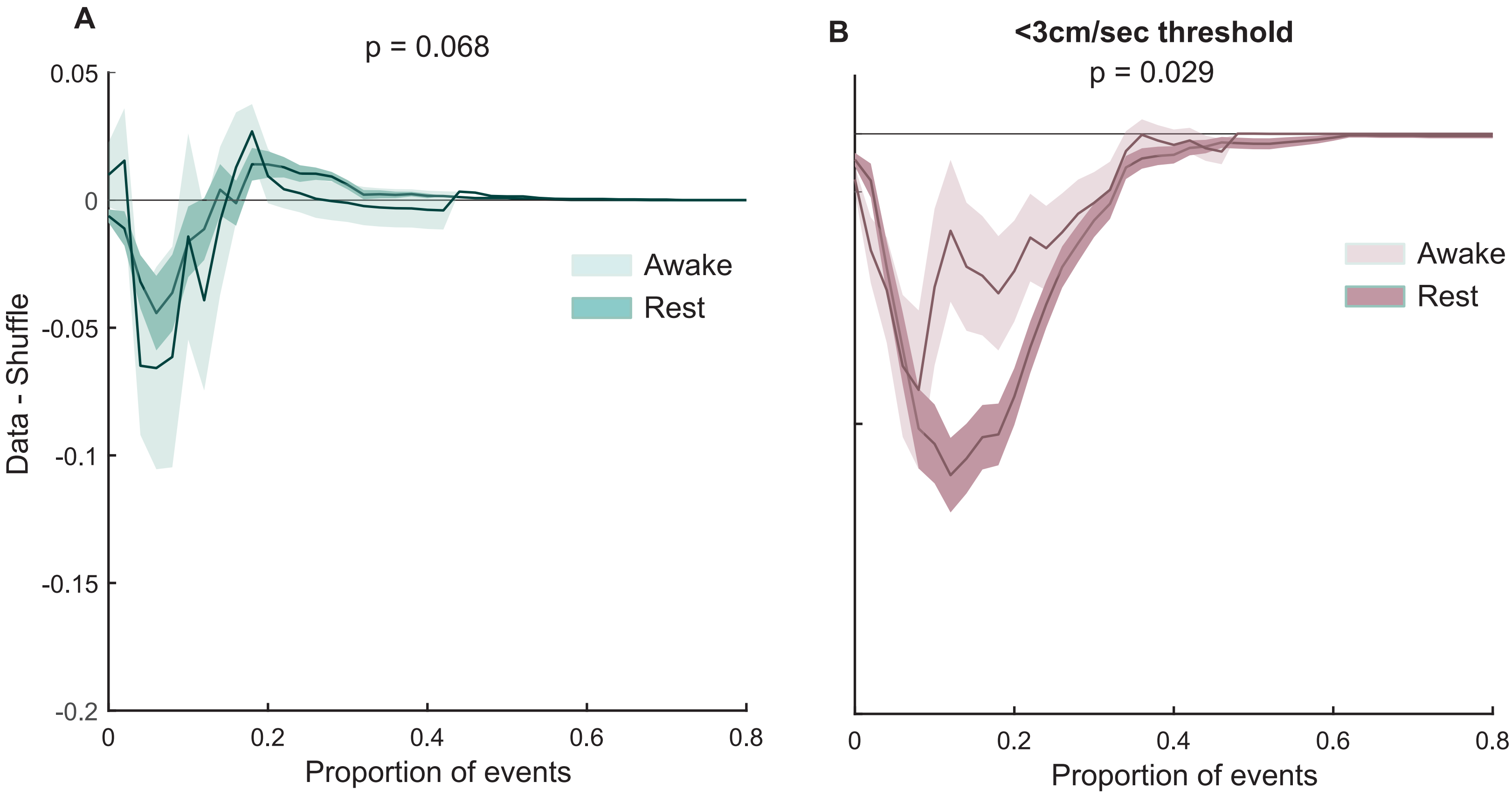


**Figure S6. Behavioural-state dependence of dMEC-hippocampal replay coordination. A**) Normalised (data-shuffle) replay coordination for non-theta modulated dMEC cells during awake (light green) and rest (dark green) periods. **B)** Normalised replay coordination for theta modulated dMEC cells during awake (light pink) and rest (dark pink) periods. Shaded are shows 1SD of bootstrapped data.
