## Supplementary material for "Theta-band phase locking during encoding leads to coordinated entorhinal-hippocampal replay": S5

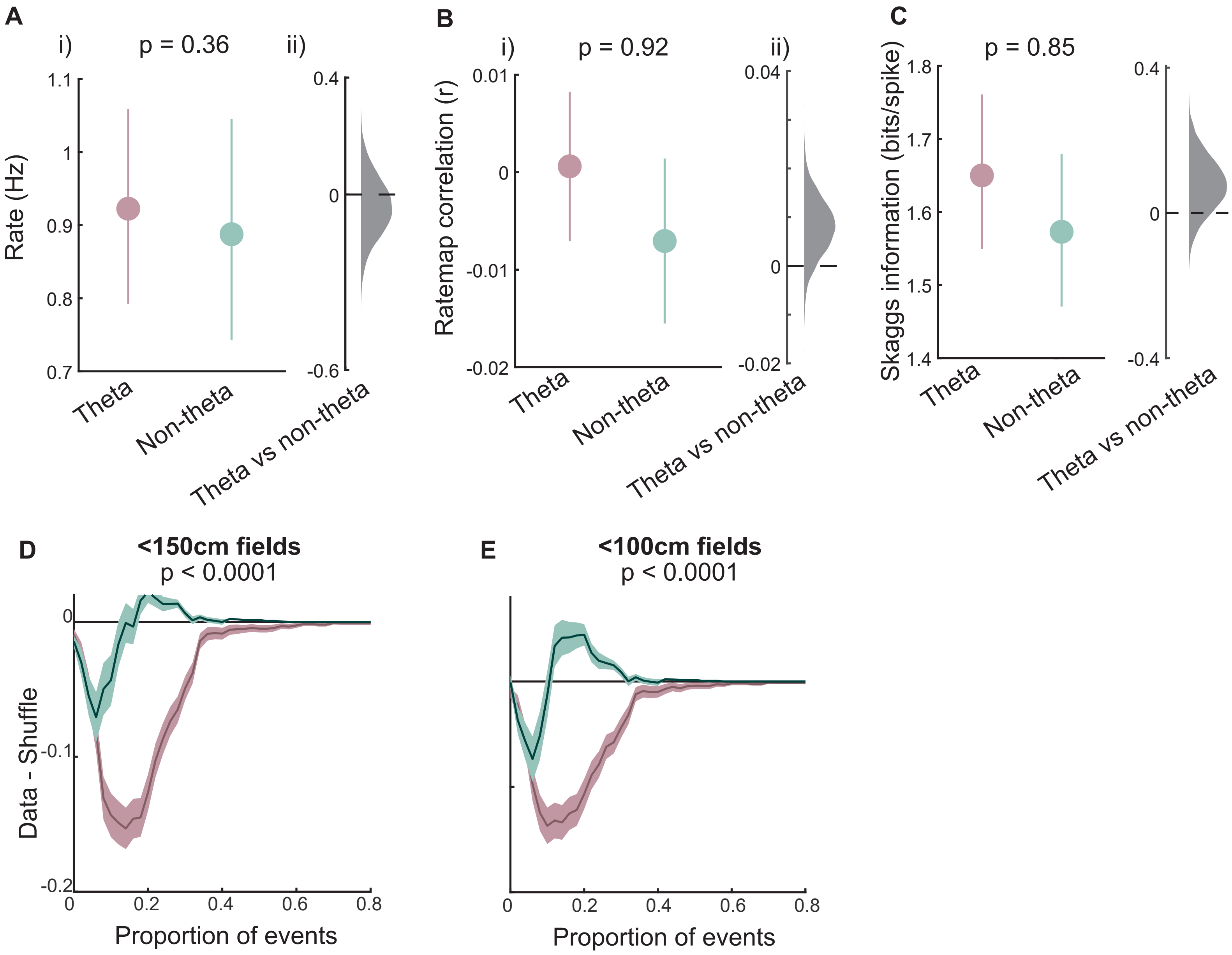


**Figure S5. Preferential coordination of theta modulated dMEC cells with hippocampal replay trajectories is not confounded by functional and activity differences between theta-modulated and non-modulated dMEC cells**. **A**) (i) Mean firing rate of theta modulated (pink) and non-theta modulated dMEC cells. Error bars show 95% CI. (ii) Kernel density of bootstrapped difference scores. **B-C**) Same as A but showing average ratemap correlations between dMEC and CA1 cells (B) and Skaggs information (C). **D**) Normalised (data-shuffle) dMEC-hippocampal replay coordination after removing dMEC cells with large (>=150cm) spatial firing fields. Shaded area shows 1SD of bootstrappe data. **E**) Same as D but using an alternative field size threshold (100cm).
