## Supplementary material for "Theta-band phase locking during encoding leads to coordinated entorhinal-hippocampal replay": S3

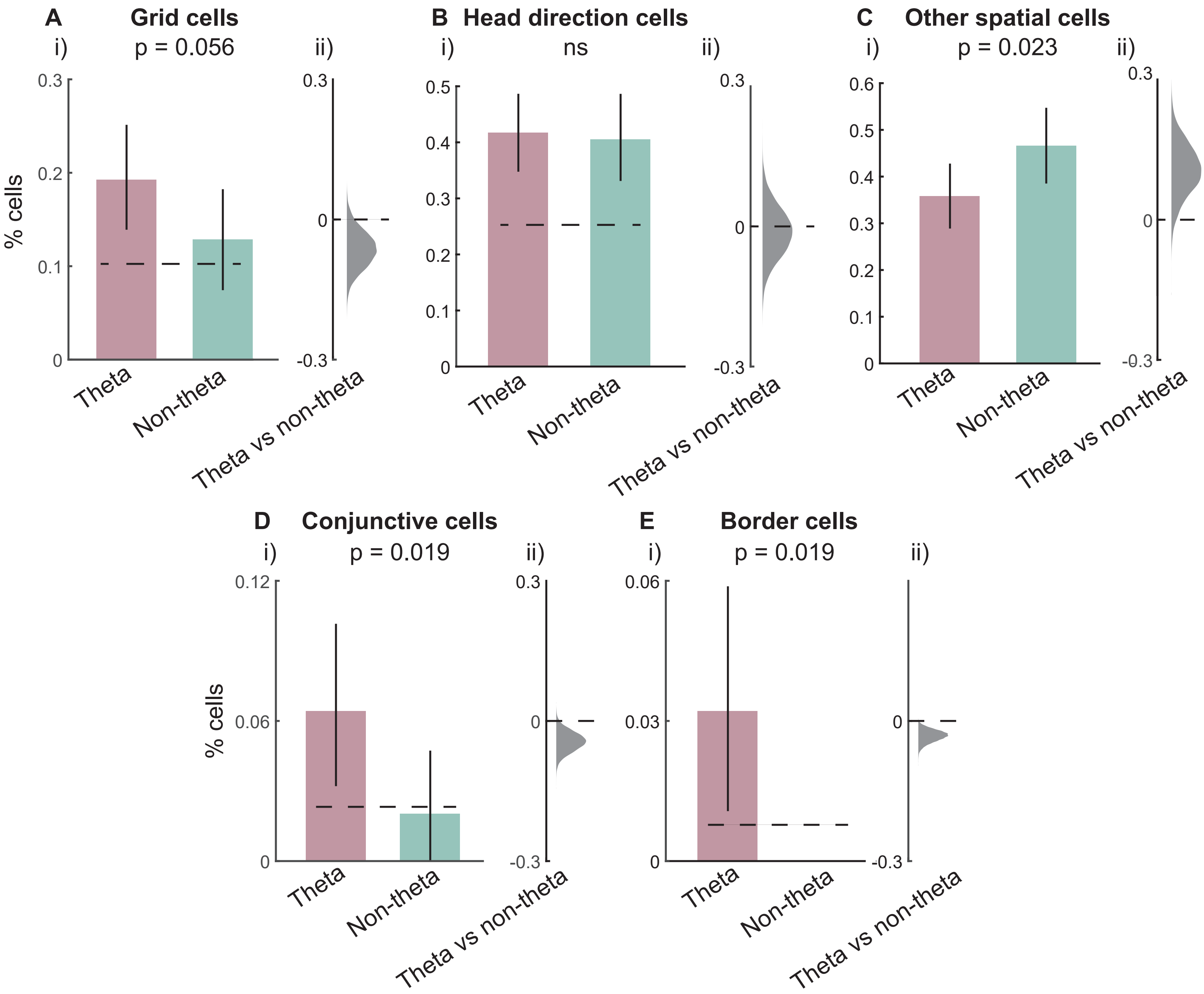


**Figure S3. Representation of distinct functional cell types in the dMEC**. **A**) i) Proportion of dMEC theta-modulated (pink) and non-modulated (green) dMEC cells that qualify as grid cells. ii) Kernel density of bootstrapped difference scores. **B-E**) Same as A but for head direction, other spatial, conjunctive and border cells.
