## Supplementary material for "Theta-band phase locking during encoding leads to coordinated entorhinal-hippocampal replay": S4

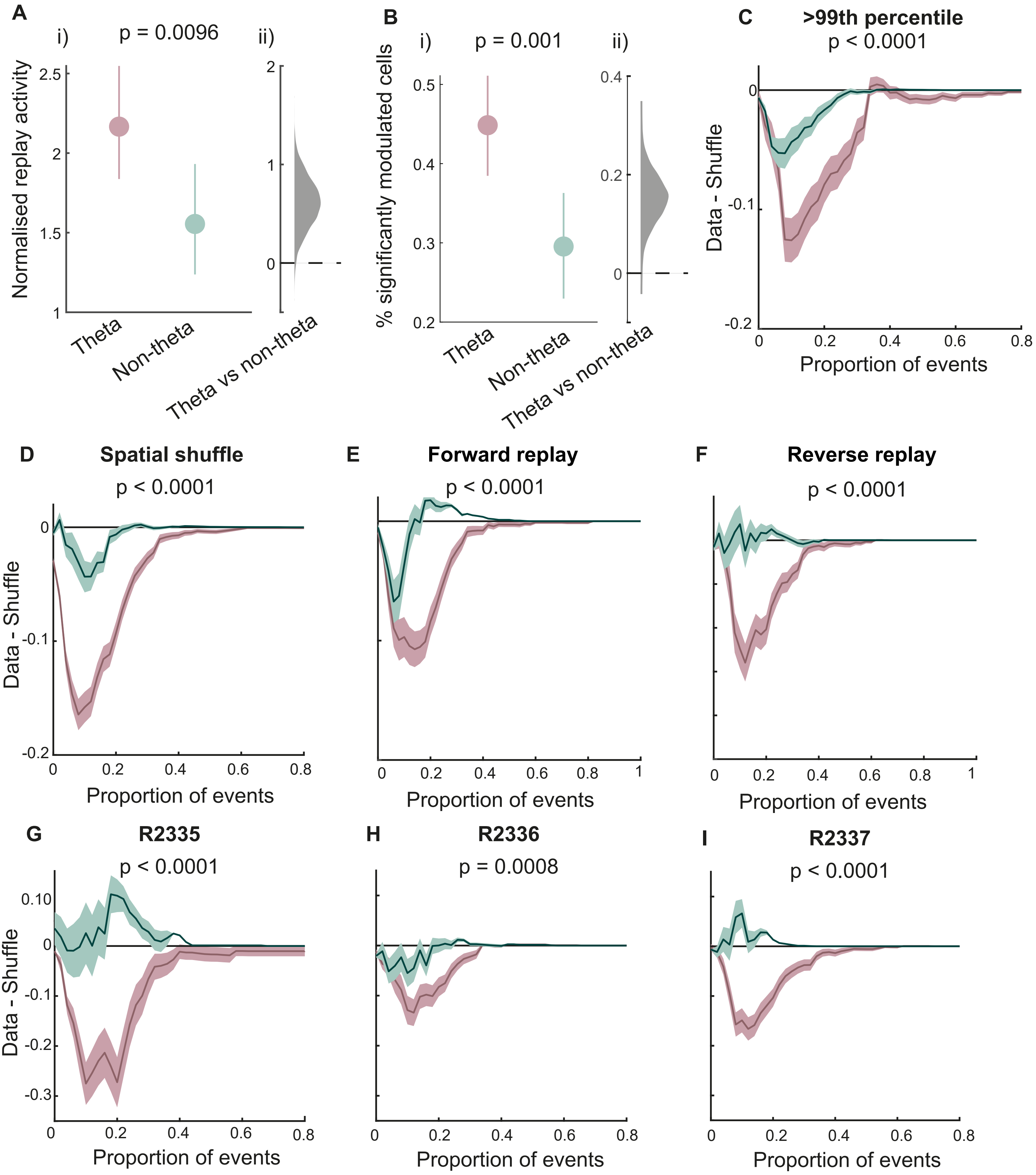


**Figure S4. Hippocampal-dMEC replay coordination: Alternative and control analysis. A**) i) Mean normalised activity modulation of dMEC theta modulated (pink) and non-modulated (green) cells. Error bars show 95% CI. ii) Kernel density of bootstrapped difference scores. **B**) i) Proportion of dMEC theta modulated and non-modulated cells that are significantly modulated by replay events, using meeasure from (A). Error bars show 95% CI. ii) Bootstrapped difference scores. **C**) Normalised (data-shuffle) replay coordination between hippocampal and theta and non-theta modulated dMEC cells using a more stringent threshold for theta modulation (99th percentile). Sahded area shows 1SD of bootstrapped data. **D-I)** Same as C but using an alternative spatial field shuffle (D), limiting analysis to forward (E) or reverse (F) replay events and repeating the analysis for individual animals (G-I).
