## Supplementary material for "Theta-band phase locking during encoding leads to coordinated entorhinal-hippocampal replay": S2

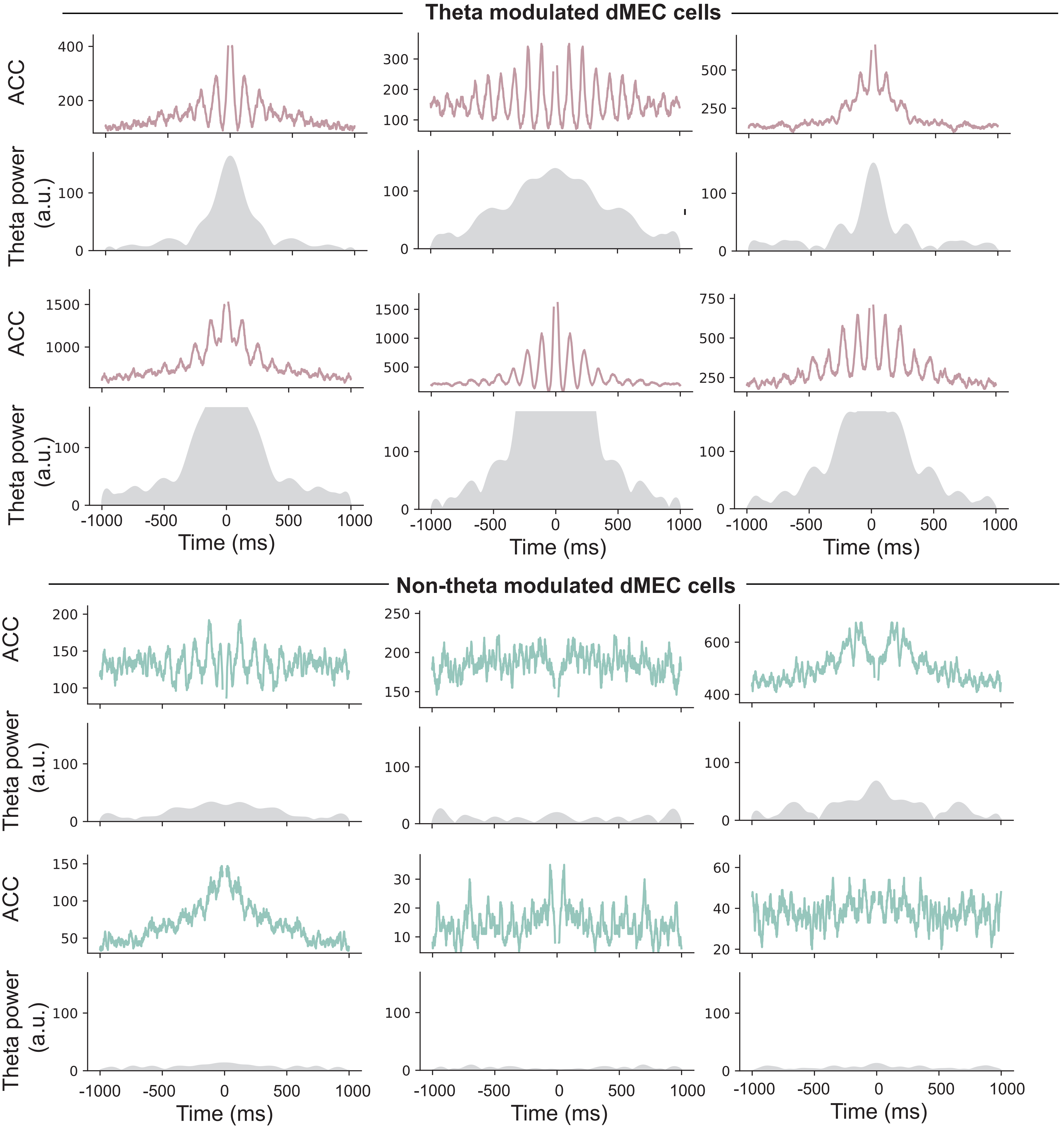


**Figure S2. Theta-band rhythmicity in dMEC cells**. Top panels: autocorrelograms for theta-modulated (pink) and non-modulated (green) dMEC cells. Bottom panel: power in the theta-band (5-12Hz) in the autocorrelogram.
