## Supplementary material for "Theta-band phase locking during encoding leads to coordinated entorhinal-hippocampal replay": S1

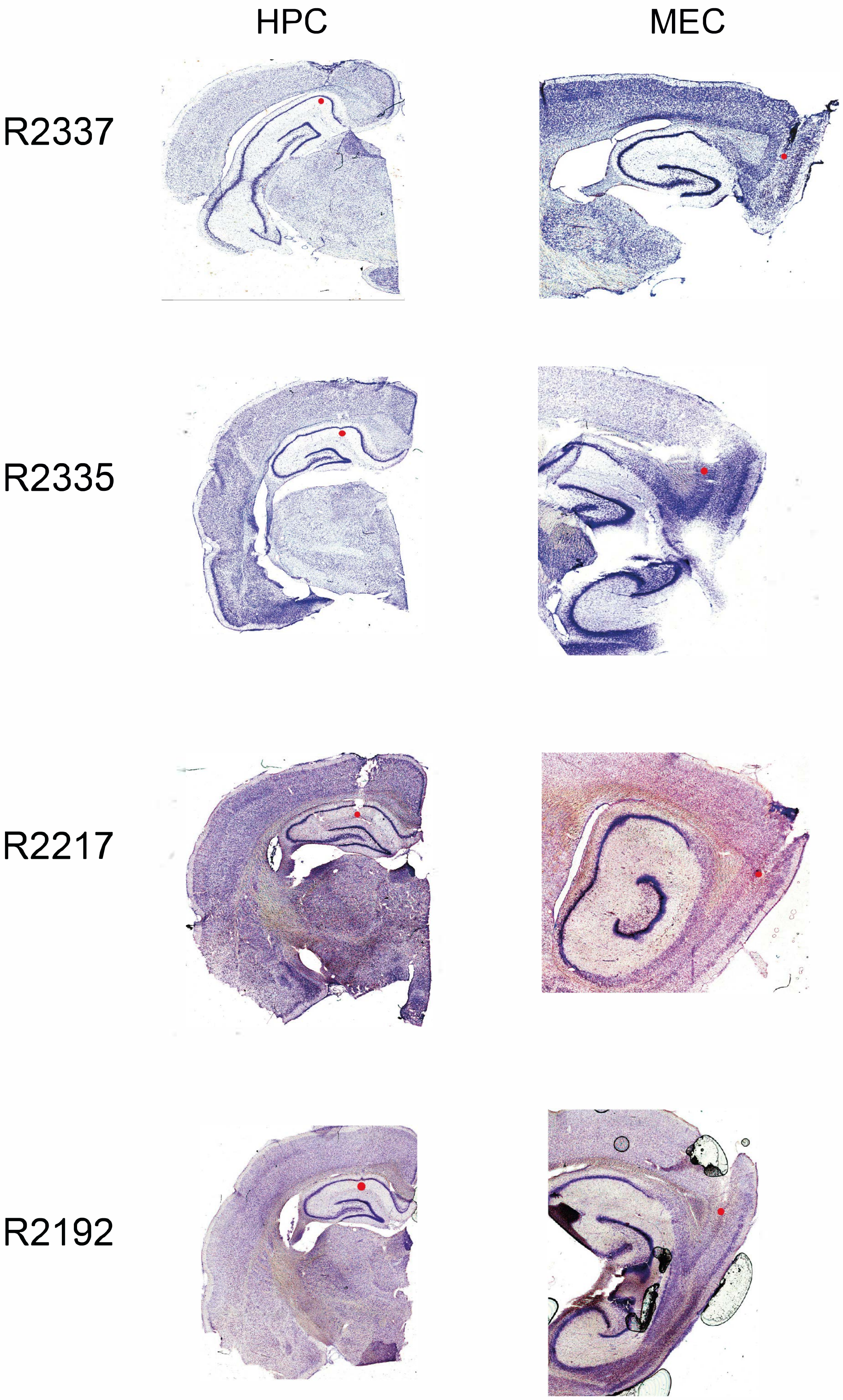


**Figure S1. Tetrode locations**. Four representative examples of Cresyl violet stained tetrode tracts from coronal (Hippocampus, left) and sagittal (MEC, right) sections. Red circle indicates the recording location for data included in this study. Left column shows rat ID.
